# Electrophysiological hallmarks for event relations and event roles in working memory

**DOI:** 10.1101/2023.05.08.539845

**Authors:** Xinchi Yu, Jialu Li, Hao Zhu, Xing Tian, Ellen Lau

## Abstract

The ability to maintain events (i.e. interactions between/among objects) in working memory is crucial for our everyday cognition, yet the format of this representation is poorly understood. The current ERP study was designed to answer two questions: How is maintaining events (e.g., the tiger hit the lion) neurally different from maintaining item coordinations (e.g., the tiger and the lion)? That is, how is the event relation (present in events but not coordinations) represented? And how is the agent, or initiator of the event encoded differently from the patient, or receiver of the event during maintenance? We used a novel picture-sentence match-across-delay approach in which the working memory representation was ‘pinged’ during the delay, in two ERP experiments with Chinese and English materials. First, we found that maintenance of events elicited a long-lasting late sustained difference in posterior-occipital electrodes relative to non-events. This effect resembled the negative slow wave reported in previous studies of working memory, suggesting that the maintenance of events in working memory may impose a higher cost compared to coordinations. Second, in order to elicit a hallmark for agent vs. patient representation in working memory, we pinged agent or patient characters during the delay. Although planned comparisons did not reveal significant differences in the ERPs elicited by the agent pings vs. patient pings, we found that the ping appeared to dampen the ongoing sustained difference, suggesting a shift from sustained activity to activity silent mechanisms. These results represent one of the uses of ERPs to elucidates the format of neural representation for events in working memory.

## 1. Introduction

The ability to represent events – not only single objects, but also the interactions between/among them – in working memory is crucial for our everyday cognition. It is clear that we can maintain ad hoc events in working memory: for example, if we see a lion hit an elephant, we are able to run subsequent mental computations that make use of the relations that compose the event representation even after the hitting itself is over (e.g., inferring that the elephant is in pain or answering a question about who was hit). However, the implementational *format* of events in working memory remains elusive.

### 1.1. The neural object indexical theory

#### 1.1.1. Evidence for the neural object indexical theory

This “format” question has become more tractable after major advances in the last 30 years in understanding how individual objects are represented and related to the local spatial context in working memory. This line of work suggests that the representation of multiple objects in working memory is supported by a limited set of indexicals or pointers also known as object files (Kahneman, Treisman & Gibbs, 1992; Pylyshyn, 1989, 2001; Xu & Chun, 2007; Carey, 2009; Zhu, Zhang & von der Heydt, 2020; Brody, 2020; Green & Quilty-Dunn, 2021; Thyer et al., 2022; Quilty-Dunn, Porot, & Mandelbaum, 2022; for a review see Yu & Lau, 2023). The number of discrete object indexicals is limited, classically suggested to have a limit of around 4 (Luck & Vogel, 1997, 2013; Cowan, 2001; Awh, Barton & Vogel, 2007; W. Zhang & Luck, 2008; Xu & Chun, 2009; Ngiam et al., 2022). Various features can be attached to these indexicals, enabling us to tell objects with some of the same features apart, and to distinguish the existence of multiple objects even when their features are identical other than spatial location (Treisman, 1998; Leslie et al., 1998). For example, if two otherwise identical flowers appear at different locations in one’s visual field, typically one is still able to distinguish this from a one-flower situation and to continue to represent the existence of both flowers in the scene even when direct visual information is interrupted by an occluding screen. This can be understood by positing the maintenance of two object indexicals in working memory, each attaching to its corresponding set of features (in this example, the two sets of features are the same except for location). In contrast, neuropsychological patients with the disorder of simultanagnosia struggle at representing two objects but not one (Coslett & Saffran, 1991; Friedman-Hill, Robertson & Treisman, 1995; Rafal, 2001), suggesting deficits related to maintaining these object indexicals.

Neuroimaging studies have suggested that these object indexicals are hosted in some subregions of the posterior parietal cortex; this being the neural object indexical theory (or the neural object file theory, Xu & Chun, 2007, 2009). This proposal is based on the observation that activities in some subregions of the posterior parietal cortex reach a plateau beyond 3-4 items (which is the classical working memory capacity), as measured by fMRI (Todd & Marois, 2004; Xu & Chun, 2006; Song & Jiang, 2006; Mitchell & Cusack, 2008; Robitaille et al., 2010; Matsuyoshi et al., 2010; Knops et al., 2014) and MEG (Robitaille et al., 2010). Consistent with the object indexical theory, these object indexicals in the posterior parietal cortex do not encode object features in themselves (Xu & Chun, 2006; Naughtin, Mattingley & Dux, 2016); rather, these indexicals connect to features represented in other regions (Xu, 2007; Naughtin, Mattingley & Dux, 2016). Taken together, evidence points to the existence of a discrete set of 3-4 object indexicals in the posterior parietal cortex, which connects to features represented elsewhere (for an illustration see Figure 1)^1^. This theory is also consistent with the proposal that the posterior parietal cortex is crucial to working memory in a more general sense (Bettencourt & Xu, 2016; K. Jia et al., 2021; for a review see Xu, 2017).

**Figure 1.**
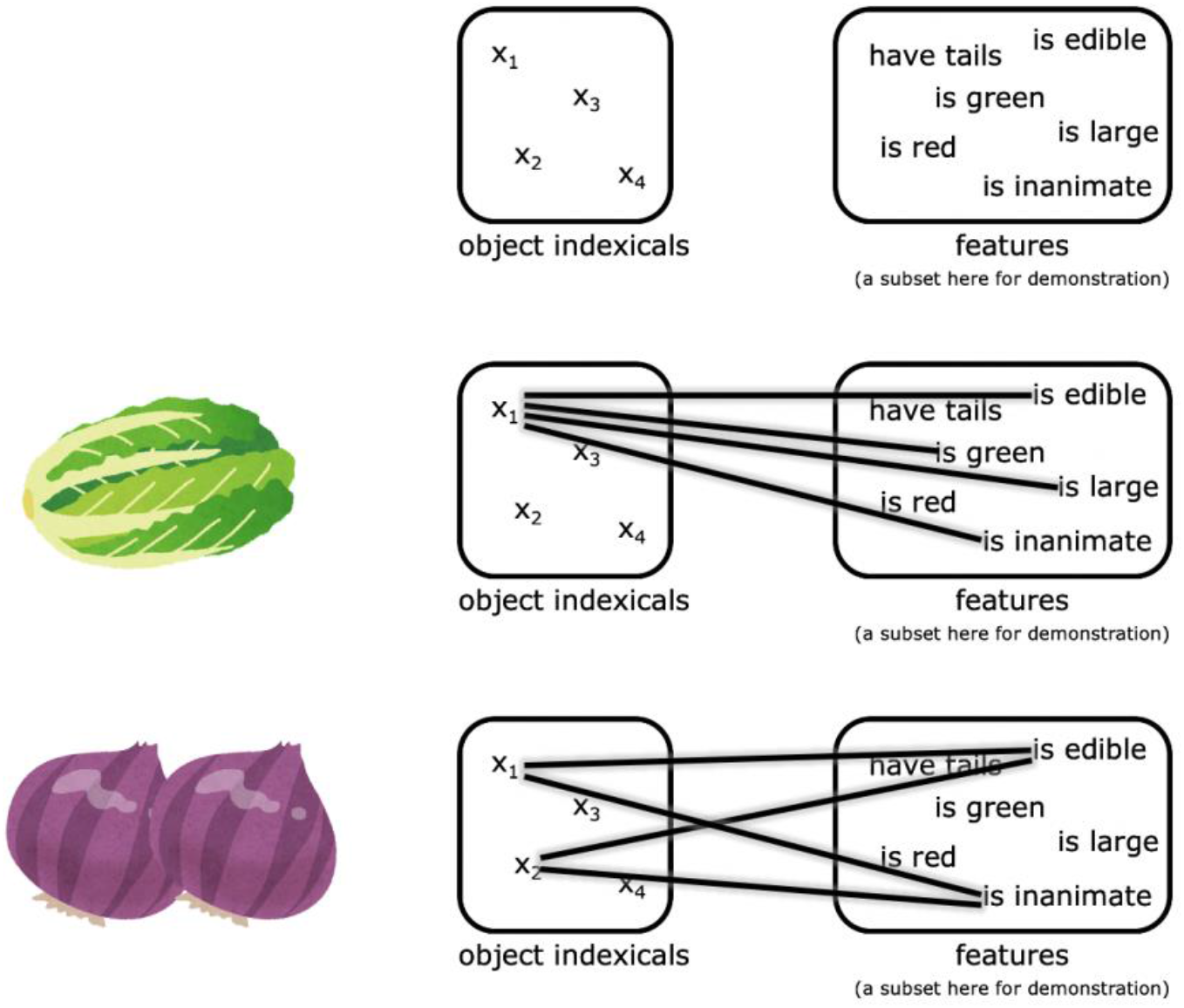
An illustration of the neural object indexical theory. The object indexicals are hosted in some subregions of the posterior parietal cortex, while the features are represented in other regions. The pictures of the Napa cabbage and onions were adapted from IRASUTOYA (https://www.irasutoya.com/), which is free for non-commercial use.

#### 1.1.2. Negative slow wave (NSW): a potential indicator for the object indexical system in working memory

Our methodology for investigating events will center around an ERP (event-related potential) component called the negative slow wave (NSW), which is thought to reflect working memory representations related to the object indexical system. Many previous ERP studies of visual working memory have observed sustained negative differences tracking working memory load for multiple objects by using a visual hemifield design that yields a response known as the contralateral delay activity (CDA) (Vogel & Machizawa, 2004). In CDA studies, multiple visual objects are presented on both sides of the visual field, but participants are asked to attend to only one side on any given trial. By hypothesis, ERPs from electrodes contralateral to the attended side will more strongly reflect the working memory representation, so corresponding posterior-occipital electrodes from each side of the scalp are subtracted from each other to compute the contralateral delay activity measure. Many studies have shown that the amplitude of the resulting CDA shows a sustained negativity that is systematically increased with an increasing number of objects to be maintained, up to the individual capacity limit of 3-4 (for an extensive review see Luria et al., 2016). Therefore the CDA has been considered as a potential hallmark for the object indexical system (e.g., Gao et al., 2013; Hakim et al., 2019; Green & Quilty-Dunn, 2021; Cai et al., 2022).

Because our current study was designed to focus on conceptual rather than visuo-spatial representation of events (i.e. the representation shared across linguistic and visual processes), we cannot expect a systematic relationship between working memory representation and visual hemifield. However, another line of visual working memory studies has demonstrated that the sustained ERP amplitude modulation by the number of maintained items can be observed even without manipulating visual hemifield. Since the measure in these studies does not depend on doing the contralateral-ipsilateral subtraction, the response is called instead the negative slow wave (NSW). The NSW has been observed in posterior-occipital electrodes with a similar time course to the CDA (Ruchkin et al., 1992; Klaver, Smid & Heize, 1999; Fukuda, Mance & Vogel, 2015; Diaz, Vogel & Awh, 2021; for a review see Feldmann-Wüstefeld, 2021)^2^. In visual delay-match-to-sample experiments where the sample objects were presented centrally or bilaterally, the NSW has been found to increase in amplitude (i.e. become more negative) as the number of items to be retained increases (Ruchkin et al., 1992; Mecklinger & Pfeifer, 1996; Klaver, Smid & Heize, 1999; Fukuda, Mance & Vogel, 2015; Feldmann-Wüstefeld, 2021; Diaz, Vogel & Awh, 2021), and reaches a plateau beyond 3-4 items (Fukuda, Mance & Vogel, 2015; L. Zhou & Thomas, 2015; Feldmann-Wüstefeld, 2021), just like the CDA^3^. Therefore, here we use the magnitude of the NSW as an index of working memory load related to the object indexical system.

### 1.2. What’s different about events? Event relations and event roles

What needs to be represented when representing events involving two objects (i.e., participants, entities; for example, “the lion hit the elephant”) in working memory, compared to simply representing two individual objects together (e.g., “the lion and the elephant”)? One is the representation of event relations (which is present in the former case but not the latter case, see Section 1.2.1 below), and the other is the representation of event roles (i.e. agents and patients, see Section 1.2.2 below).

Note that in the current article, our major interest lies in the *conceptual* representations that are shared across visual and linguistic processes (cf. Jackendoff, 1987; Wurm & Caramazza, 2019; Fitch, 2020; Mahon & Kemmerer, 2020). That is, we are interested in the representation that is shared both when one is viewing the scene where a lion hits an elephant, and/or when one reads a sentence like “the lion hit the elephant”.

#### 1.2.1. The neural representation of event relations

By event relation, we refer to an intuitive distinction between situations where an elephant hits/points at/pushes a lion vs. an elephant and a lion simply standing beside each other and not doing anything to each other (see Figure 2). The difference between these two kinds of situations is whether there is an event relation. Of course, there are different *types* of event relations (i.e., event types), e.g., hitting, pointing at, pushing; but the focus of the current article is how our neural system responds to the existence of an event relation at all in working memory, not how we represent different event types.

**Figure 2.**
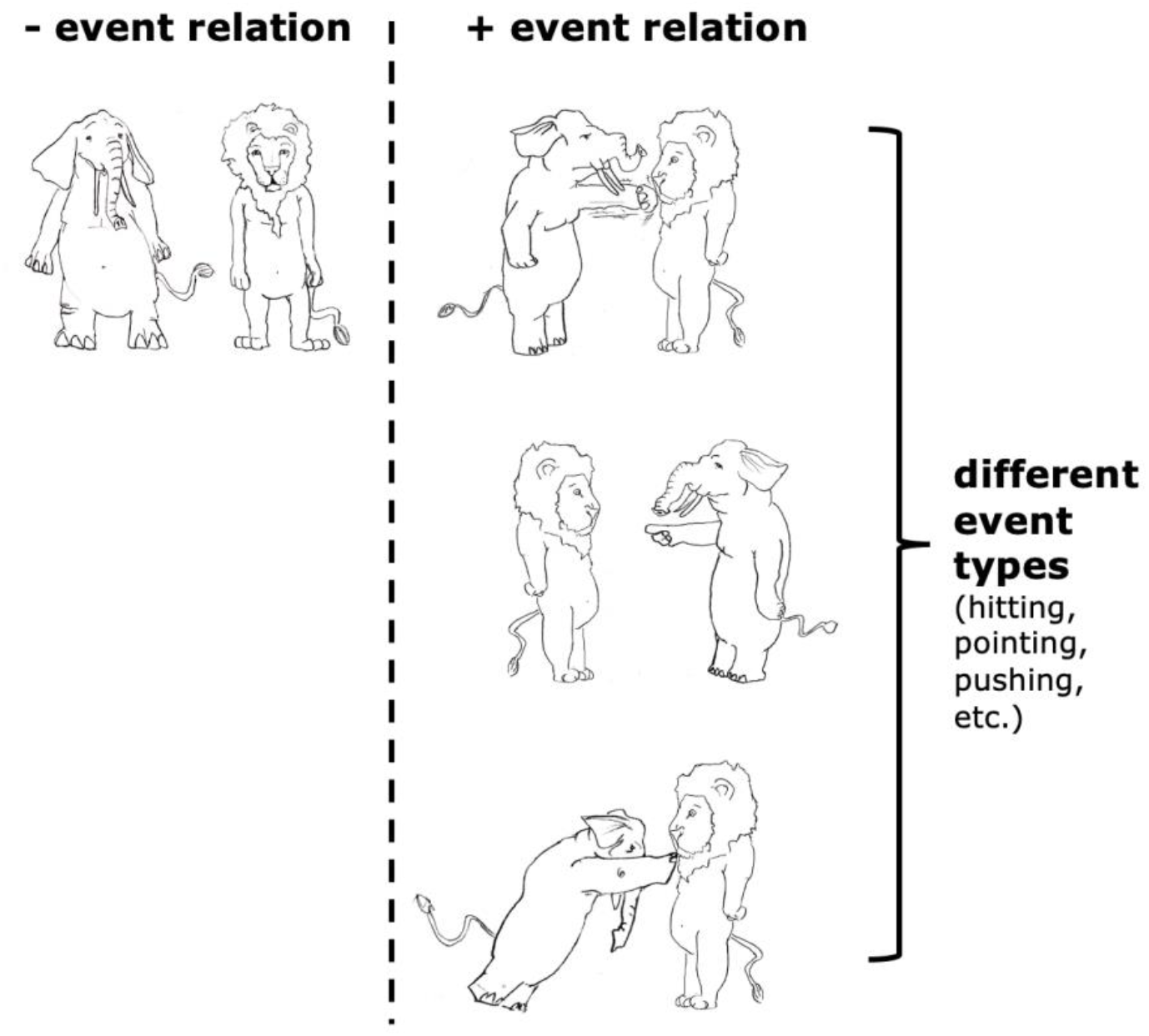
An illustration of “event relations” that we refer to in the current article, in contrast to “event types” (i.e., “the types of event relations”). In the situations depicted in the right column, there are event relations, yet in the left column there isn’t.

##### The extraction of event relations

The existence of event relations can be extracted from both visual and linguistic stimuli. For example, we can tell whether there is an event relation for each situation in Figure 2; we can also tell whether there is an event relation from sentences like “The lion and the elephant stood there” vs. “The lion hit the elephant”.

In terms of visual studies, although many studies have addressed the speed of extracting the event type (Dobel et al., 2007; Hafri, Papafragou & Trueswell, 2013; de la Rosa et al., 2015), less is known about how fast the brain detects whether there is an event relation at all; the only study we are aware of is Hultén et al. (2014). Using MEG, Hultén and colleagues observed that visual events (e.g., where a lion hits an elephant), compared to visual coordination (e.g., where a lion and an elephant were simply standing beside each other), triggered an early (∼ 200 ms) difference in left occipital regions. The effect observed in Hultén et al. (2014), based on its location (i.e. occipital regions), is likely visual rather than conceptual in nature.

As for linguistic studies, Gaston (2020) investigated the difference between linguistic stimuli including event relations compared to coordination using MEG. Gaston (2020, Experiment 4) examined the brain responses to serially presented, nonsensical linguistic input (e.g., the | toasty | tractors | and *or* entered | the | scenic | cathedrals) and observed a difference from ∼ 150 ms to ∼ 550 ms after the onset of “and” vs. a verb in regions including the transverse temporal sulcus, pSTS (posterior superior temporal sulcus), STS, and ATL (anterior temporal lobe). This effect is in line with the observation of Matchin et al. (2019a), that the presentation of a verb within a sentence elicits a difference starting from ∼ 450 ms upon onset in the angular gyrus, compared to a baseline consisting of phrases without a verb. Such transient effects may reflect a difference in linguistic structure building and/or the representation of event relations at the conceptual level. We should note that the analysis pipelines of Gaston (2020) and Matchin et al. (2019a) may not be sensitive to long-lasting sustained effects.

##### The maintenance of event relations in working memory

To our knowledge, only two behavioral studies have directly addressed this question (Clevenger & Hummel, 2014; Shen et al., 2021). The results of Clevenger & Hummel (2014) suggested that more relations impose higher working memory load, and the results of Shen et al. (2021) suggested that only two event relations can be maintained in working memory. However, these results remain inconclusive in terms of the *format* of the representation of event relations in working memory; specifically, how are event relations represented with respect to the object indexical system?

One way to approach this question, and the one we pursue in the current study, is to compare the maintenance of events like “the tiger hit the elephant” and the maintenance of non-events like “the tiger and the elephant”, which we will refer to as “coordinations”. To our knowledge, the MEG study from Hultén et al. (2014) is the only previous neurophysiological study to have examined the maintenance of events compared to coordinations^4^. In their study, pictures of events and coordinations were presented for 1500 ms, followed by a speech production task. A numerical difference was observed between events and coordination conditions in a somewhat late time window (from ∼ 300 ms to at least 1 s) time-locked to picture onset in the posterior parietal cortex. Although this difference did not survive statistical significance in their analysis, this could be due to the small cohort of subjects (N=10).

How should we expect event representation in working memory to manifest differently from simply representing a pair of objects? Through the lens of the object indexical system, we think that there are at least two major possibilities (see Figure 3). (a) The event relation may be represented with a separate object indexical from the participants (cf. Clevenger & Hummel, 2014; Shen et al., 2021). In other words, adding an event relation increases working memory cost in exactly the same way as adding another object, and counts against the same capacity limit as objects. For example, in the case of “the tiger hit the elephant”, apart from the two object indexicals for the tiger and the elephant, there is a separate object indexical representing the hitting relation, possibly by connecting to the concept HITTING and other features of hitting. In this case, there will be an NSW-like difference between the maintenance of events and coordinations. We want to especially highlight that in this hypothesis, just as the indexicals for objects are not representing the features of the objects in themselves (but connecting to those features elsewhere), the indexicals for event relations are not representing the event types in themselves (but connecting to relevant features and knowledge elsewhere) too. (b) Another possibility is that the object indexical system in working memory does not in itself explicitly encode event relations between objects, as implied in G. Altmann & Ekves (2019). The relational encoding might be accomplished by other posterior parietal circuits, or by other brain regions altogether, such as the hippocampus (e.g., Konkel & Cohen, 2009; Eichenbaum & Cohen, 2014). The object indexical working memory representation might only code such relations implicitly in an object-centered fashion, for example by connecting agent and patient “features” to the object indexicals (for further explanation of agents and patients see Section 1.2.2 below). In this case, there is no reason to predict that there will be an NSW-like difference between the maintenance of events and coordinations.

**Figure 3.**
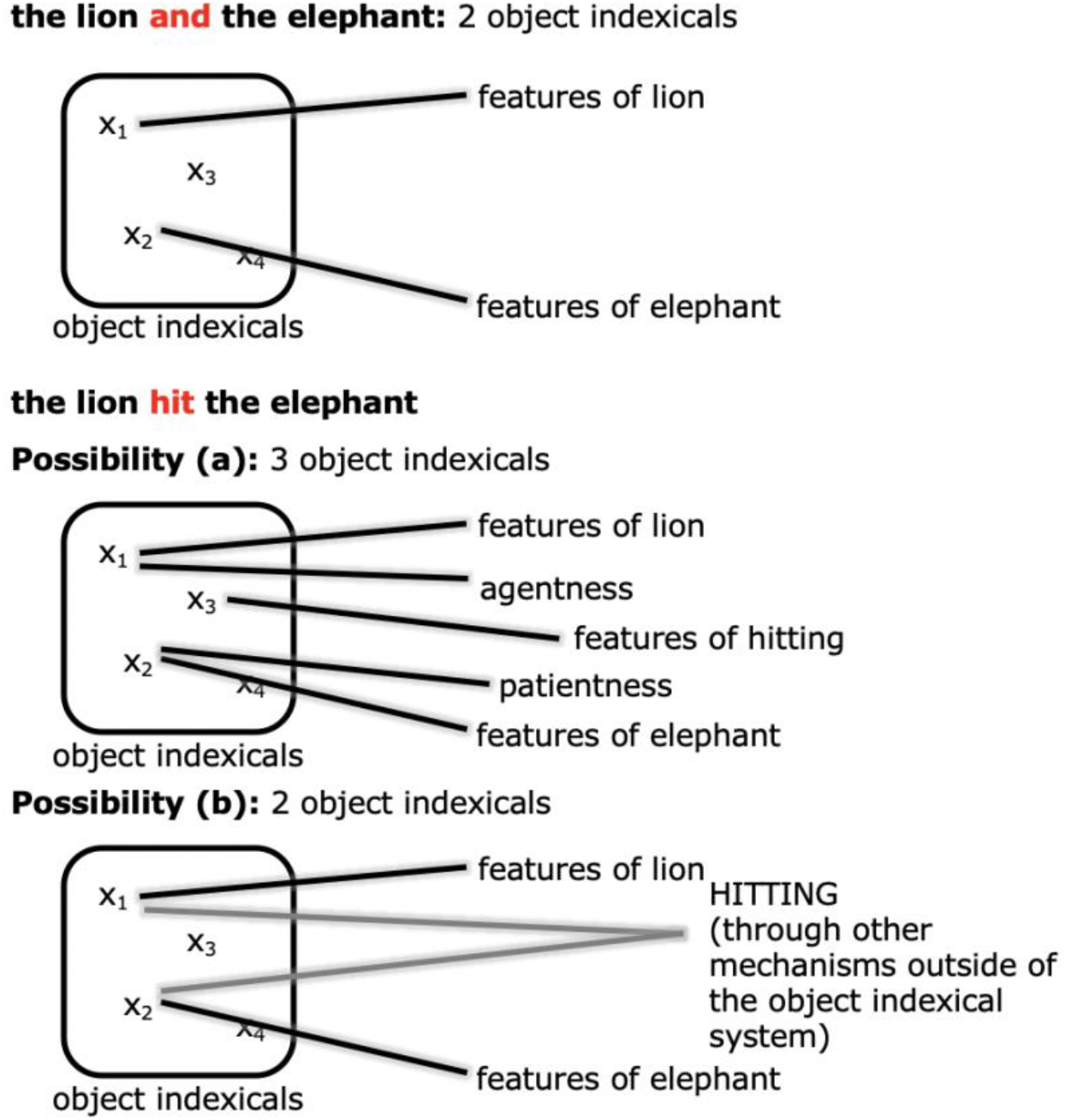
An illustration of the two possibilities of how event relations are represented. Possibility a: the event relation is encoded by another object indexical. Possibility b: the event relation is represented by other mechanisms in e.g., other regions of the posterior parietal cortex or the hippocampus.

#### 1.2.2. The neural representation of event roles

Many modern theories assume the existence of generalized roles like agent (the initiator of the event) and patient (the receiver of the event), which characterize the involvement of participants across many different types of events (see Strickland, 2017; Rissman & Majid, 2019; Ünal, Ji & Papafragou, 2021; Wilson, Zuberbühler & Bickel, 2022). This proposal roots especially from the tradition in the field of formal semantics (for overviews see Williams, 2015, 2021), which may be capturing these roles at the linguistic semantic level or in fact at the non-linguistic conceptual level. How such non-linguistic event roles (also known as “semantic” roles or participant roles) are neurally coded in working memory remains unclear.

##### The extraction of event roles

Conceptual event roles can be extracted from visual and linguistic stimuli. In visual studies (i.e., studies using visual stimuli), it has been observed that the event role of objects can be extracted after a brief visual presentation (Dobel et al., 2007; Hafri, Papafragou & Trueswell, 2013). To our knowledge, there has been only one neurophysiological study that directly addressed the encoding of event roles from visual presentations (Cohn, Paczynski & Kutas, 2017). In their EEG study, subjects were presented with comic strips (modified from *the Peanuts*) sequentially, and the ERP responses to frames illustrating the to-be-agent and to-be-patient character were compared. Of the to-be-agent characters, some were making actions suggesting their agentness (“preparatory agents”), while some weren’t (“non-preparatory agents”). Comparing the preparatory agent condition with the other two conditions (i.e. the non-preparatory agent condition and the patient condition), a left anterior negativity in 300-600 ms and a slightly leftward fronto-central positivity in 500-900 ms were identified. This may reflect the difference between extracting the agent object vs. the patient object, or in fact the difference between extracting the agent object vs. an object with a not-yet-determined role.

One of the ways that some languages convey event role assignments is by linguistic case markers. For example, in Japanese, the conceptual content of *John gave Mary a book* can be realized linguistically as “John-NOM Mary-DAT book-ACC gave”, where -NOM, -DAT, -ACC stand for nominative, dative and accusative case markers *ga*, *ni*, and *o*. Apart from case markers, language can convey event role assignments in other ways such as word order / construction / structural position; for example, in both “The cat chased the dog” and “The dog was chased by the cat”, we know that the cat is the agent and the dog is the patient. Attempts have been made using fMRI to decode the binding between event roles and the type of corresponding object (e.g., cat or dog) using this kind of English sentences (e.g., Frankland & Greene, 2015, 2020; J. Wang et al., 2016; Lalisse & Smolensky, 2021), yet due to the poor temporal resolution of fMRI, these studies were not optimal for disentangling the extraction and maintenance processes of event roles.

##### The maintenance of event roles

Behaviorally, it has been observed that maintaining more event role-object bindings reduces the accuracy of subsequent memory performance (Jessop & Chang, 2020), yet it is unclear whether this burden was driven by the increase in the number of objects held in working memory or the increase in the number of event roles bound to these objects (or both). It has also been found that representing conflicting event roles (e.g., being the agent in one event but the patient in another event) compromises memory performance (Jessop & Chang, 2022). However, we are still far from a complete understanding of these behavioral observations. Notably, Peng et al. (2017), through a behavioral experiment using the dense sampling method, suggested that event roles are represented at (or even by) different phases of the alpha band (∼ 10 Hz) rhythm (for further discussions of this experiment see Section 3.2).

To our knowledge, no prior electrophysiological study has examined the maintenance of event roles in working memory. In fact, it is a practical challenge to disentangle the representation of the agent object and the patient object in working memory, given that they are in the same event. In the current study we introduce an exploratory paradigm adapted from the “pinging” paradigm used in recent working memory studies (Wolff et al., 2015, 2017, 2019, 2020; Fornaciai & Park, 2020; ten Oever, De Weerd & Sack, 2020; Huang, H. Zhang & Luo, 2021; Y. Fan & Luo, 2022; Duncan, van Moorselaar & Theeuwes, 2023). Wolff et al. (2015) conducted a standard delay-match-to-sample working memory task in which the match in the orientation of two Gabor patches needed to be evaluated across a relatively long delay. During the delay period, a high-contrast visual bull’s-eye-shaped stimulus with no orientation information (i.e. the ping) was presented in an attempt to probe the encoding of the orientation information being maintained in memory, similar to the idea of sonar or echolocation. It was found that the otherwise undecodable memory content at that time (the orientation of the sample Gabor patch in this case) was decodable if the ping was presented. In this study and others that followed, the main idea was to probe different working memory contents during retention with the same impulse perturbation (i.e. the ping).

In the current study, we adapt the pinging paradigm to probe different parts of the same working memory content during retention using different pings (see also Y. Fan et al., 2021). Essentially the idea for our current experiment (see Figure 4) is that the event roles are instantiated by an initial image of the event, and while being carried forward in working memory we “ping” the event role with the name of the entity to which it was bound (e.g. the word “lion” pings the agent in this example, thus being an “agent ping”). We ping the agent in half of the event trials and the patient in the other half, and therefore we are able to compare the ERP corresponding to agents and patients in working memory directly. Notably, since each animal can be the agent or patient, the agent pings and patient pings are perceptually the same. In order to keep the trial structure consistent throughout the experiment, we also pinged one of the two animals in the coordination condition.

**Figure 4.**
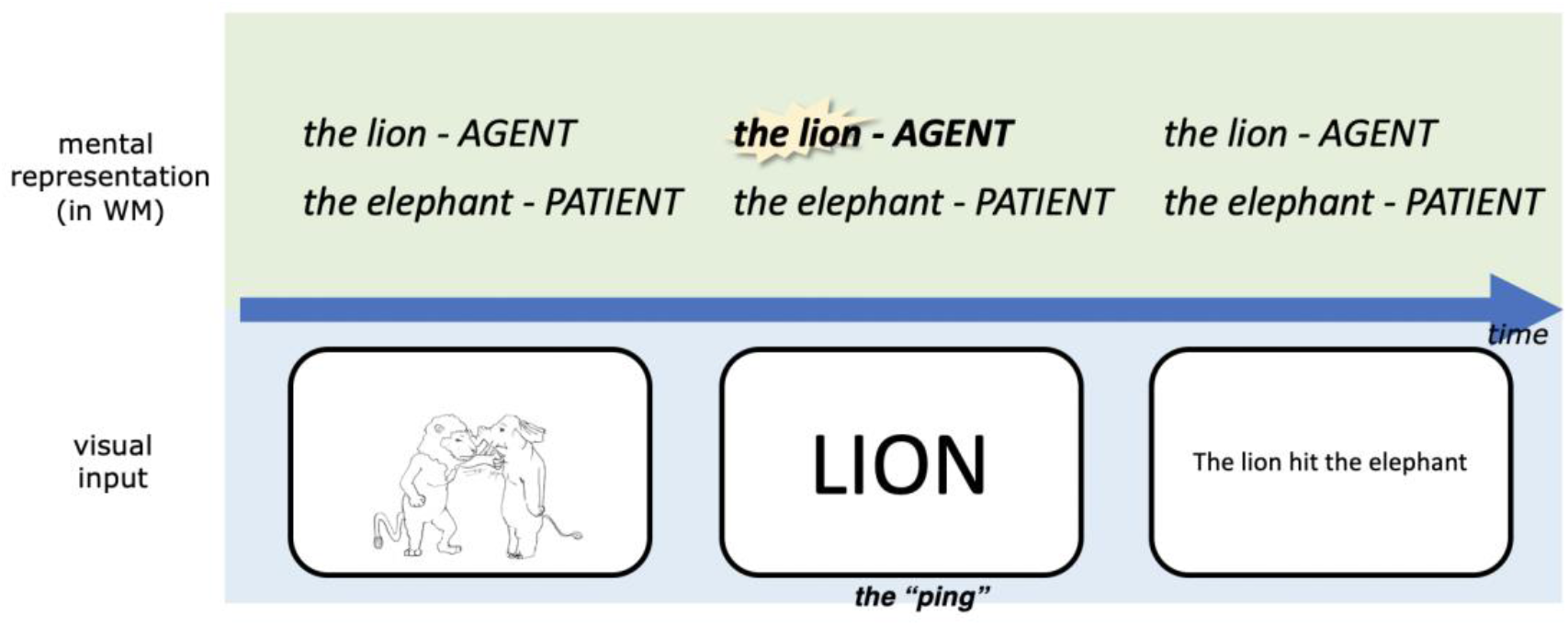
A simplified schematic illustration of our experimental design, highlighting our adapted pinging paradigm. The task was to respond to whether the linguistic expression matched with the picture in this trial upon presentation of the linguistic expression, this was designed to encourage subjects to represent the event *conceptually* during the retention period. WM: working memory. For detailed parameters of the experiment, see Materials and methods.

Although it is as yet unclear how event roles are represented, one possibility that we explored in this study is that event roles are represented as a kind of “magnitude”^5^. Some authors have proposed that regions of parietal cortex represent diverse kinds of scalar and quantity information in a common neural implementation format (Walsh, 2003; Bueti & Walsh, 2009; Summerfield, Luyckx & Sheahan, 2020; for a critical review see B. Martin, Wiener & van Wassenhove, 2017). It has been found that magnitudes (including numerosity, magnitude of reward, and magnitude of evidence during decision-making) drive an ERP effect in central-parietal electrodes, that is, the central parietal positivity (CPP, Yeung & Sanfey, 2004; O’Connell, Dockree & Kelly, 2012; Spitzer, Waschke & Summerfield, 2017; Luyckx et al., 2019; for an overview see O’Connell & Kelly, 2021). For transient (i.e. not constantly-changing) stimuli, it is a late component at around 300 to 700 ms post stimulus-onset (Spitzer, Waschke & Summerfield, 2017; Luyckx et al., 2019), with larger magnitudes being more positive. Behaviorally, agent-patient event roles have sometimes been observed to interact with spatial position (i.e. left-right), which can be treated as a “magnitude” (e.g., Chatterjee, Maher & Heilman, 1995; Maass & Russo, 2003; Dobel, Diesendruck & Bölte, 2007; but see Geminiani et al., 1995; Barrett et al., 2002; L. J. Altmann et al., 2006; Kazandjian et al., 2011; Dobel et al., 2014 for conflicting results). The hypothesis that event roles are represented as magnitudes would predict an ERP effect with a similar scalp distribution to the CPP.

We ran two experiments with a similar design in two sites, one in Chinese in China (Experiment 1) and one in English in the US (Experiment 2). We take posterior-occipital electrodes P7, P8 as the NSW region of interest (ROI), and central-parietal electrodes CP1, CP2, Pz (Experiment 1) and Cz, CPz, Pz (Experiment 2) as the CPP ROI (see Figure 5).

**Figure 5.**
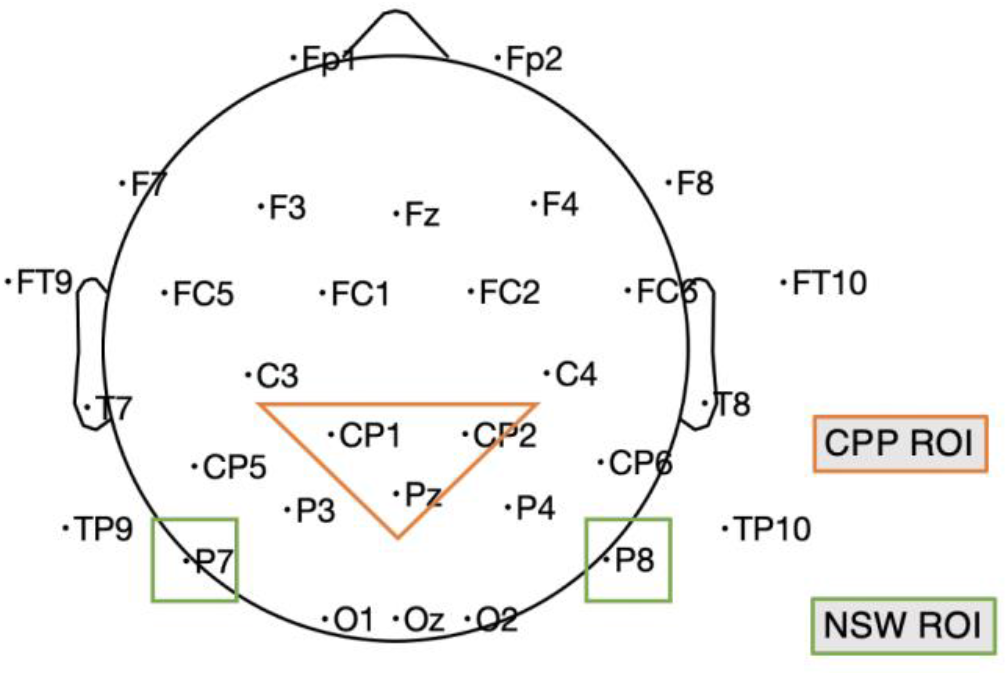
Illustration of the electrodes of our two ROIs in Experiment 1; in Experiment 2 the NSW ROI included Cz, CPz and Pz because of a different layout. CPP: centro-parietal positivity; NSW: negative slow wave; ROI: region of interest. This is based on the standard 10-20 map provided by the DIPFIT plug-in of EEGLAB.

## 2. Results

### 2.1. Experiment 1

#### 2.1.1. Behavioral results

Across the 19 subjects included in the analyses, the mean proportion of correct responses was 96% (range 89%-99%). Mean proportion of correct responses was 93% (when the agent was pinged), 95% (when the patient was pinged) and 98% (in coordination trials). Based on the high accuracy, all trials were analyzed for the EEG analysis.

The mean (± SD) reaction time of correct responses was 1241 ± 288 ms when the agent was pinged, 1230 ± 275 ms when the patient was pinged, 1236 ± 278 ms for all event trials and 903 ± 233 ms for all coordination trials. The reaction times reported here were first averaged within subjects for all the correct trials, then averaged across subjects. Paired t-test (two-tailed) indicated no significant difference between the reaction times for agent ping trials and patient ping trials, *t*(18)=0.6, *p*=0.56. Paired t-test (two-tailed) revealed a significant difference between the reaction times for event trials and coordination trials, *t*(18)=11.7, *p*<0.001. It has been observed with the delay-match-to-sample paradigm that the more objects one needs to maintain in working memory, the slower one reacts to the probe stimuli (Hyun et al., 2009; Mitchell & Cusack, 2011; Park, W. Zhang & Hyun, 2017). Therefore, the longer reaction time to the probe (in our experiment the linguistic expression) for event trials observed in our current experiment is in line with our interpretation of our EEG results, that the maintenance of events imposes a higher working memory load compared to the maintenance of coordinations, similar to representing an extra object (see Section 3).

#### 2.1.2. EEG results

ERP responses to event pictures and coordination pictures are illustrated in Figure 6. We expected initial evoked responses to the event pictures and coordination pictures to differ because the pictures in the two conditions were physically different across various dimensions, and that responses to the sentences in the two conditions might differ because the form of the correct response differed in the two conditions. Our question of interest was whether we would observe differences across the delay between the picture presentation and the sentence presentation in the posterior electrodes associated with the negative slow wave.

**Figure 6.**
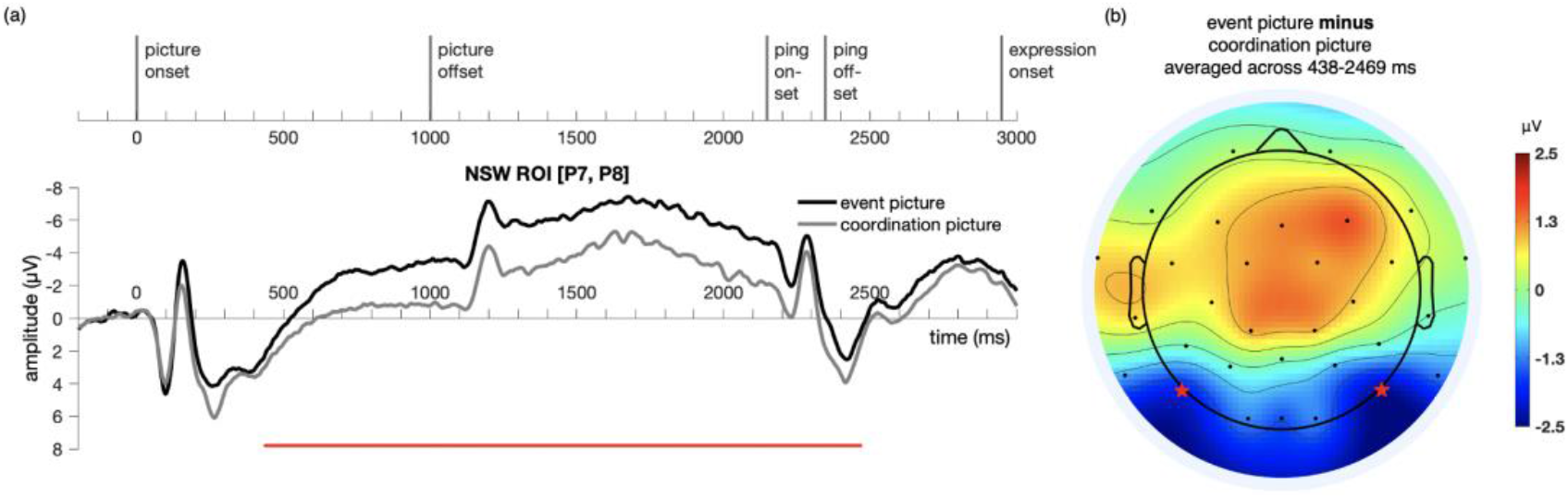
Illustration of the results for epochs time-locked to picture onset in the NSW ROI (electrodes P7 and P8, as marked by the stars in Figure b) for Experiment 1. (a) The significant cluster (438-2469 ms) is marked in red. (b) The scalp distribution average across the 438-2469 ms time window. NSW: negative slow wave; ROI: region of interest.

A temporal cluster-based permutation test (Maris & Oostenveld, 2007) was performed on the responses to the event picture and the coordination picture in the time window of -200:3500 ms for the NSW ROI using customized MATLAB scripts. First, potential clusters were identified with point-to-point paired t-test at a threshold of alpha = 0.05 (two-tailed). Then, the t values of these potential clusters were compared against a permuted (permutations = 9999) distribution of t values, with a threshold of alpha = 0.05 (two-tailed). A cluster was identified in the negative slow wave (NSW) ROI, extending from 438 to 2469 ms after picture onset (*p*=1×10^-4^, Figure 6). The difference manifested as an increased negativity for the event pictures relative to the coordination pictures in these posterior electrodes. Although our analyses focused on the pre-determined NSW ROI, the scalp topography of the cluster suggests that this posterior negativity may have been accompanied by a corresponding (central-)frontal positivity.

ERP responses to the delay-period agent ping and the patient ping in the event condition are illustrated in Figure 6. Cluster-based permutation test (Maris & Oostenveld, 2007) was performed comparing the agent ping vs. patient ping conditions in the time window of -200:1800 ms using customized MATLAB scripts using the same approach described above for the event picture comparison. No significant clusters distinguishing the agent ping and patient ping were identified in the CPP ROI, nor in the NSW ROI (Figure 7).

**Figure 7.**
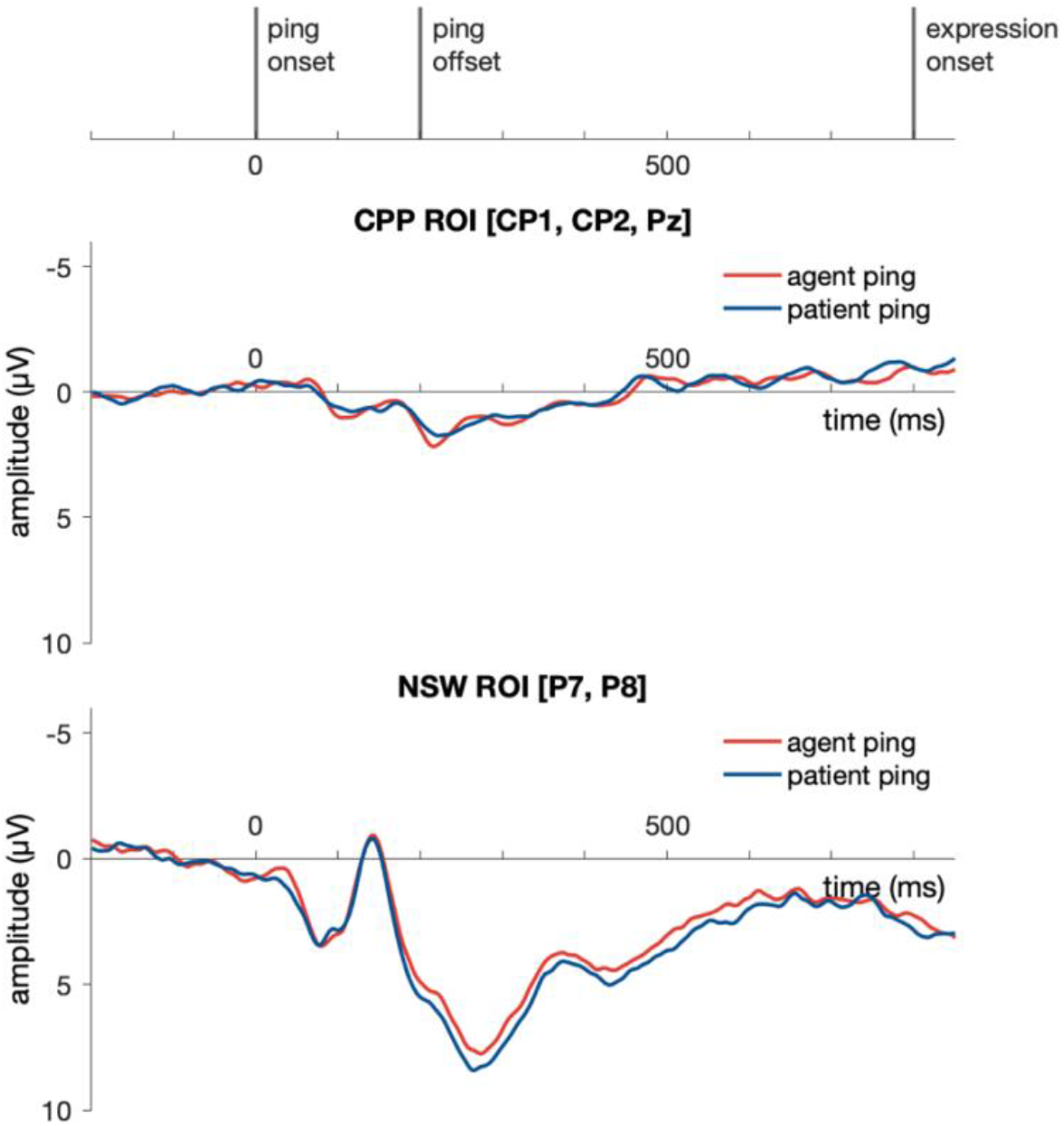
(Left) Illustration of the results for epochs time-locked to ping onset in the NSW ROI and the CPP ROI for Experiment 1. NSW: negative slow wave; CPP: centro-parietal positivity; ROI: region of interest.

#### 2.1.3. Rationale for Experiment 2

In Experiment 1, we observed that maintaining events in working memory, compared to maintaining coordinations, accompanied a sustained NSW (negative slow wave) effect. However, during data analysis we discovered an issue in the accuracy of the timing parameters that could have led us to overestimate the duration of the sustained negativity (see Section 4.1.6 for more detail). In Experiment 2, we aimed to replicate the effect in a different population (Experiment 1 was run in Shanghai with stimuli in Chinese and Experiment 2 was run in the US with stimuli in English), with corrected presentation parameters that allow more precise estimation of the duration of the effect.

As visual inspection of the Experiment 1 effects suggested that the presentation of the ping may have acted to reduce or eliminate the sustained negativity, we also took Experiment 2 as an opportunity to evaluate this possibility by presenting the ping somewhat earlier in the trial. We presented the pictures for shorter duration in Experiment 2 (400 ms) compared to Experiment 1 (1000 ms), as an additional measure to ensure that the sustained effects we observed did indeed reflect delay-period activity rather than continued visual processing.

### 2.2. Experiment 2

#### 2.2.1. Behavioral results

Across the 16 subjects included in the analyses, the mean proportion of correct responses was 92% (range 83%-97%). Mean proportion of correct responses was 88% (when the agent was pinged), 91% (when the patient was pinged) and 94% (in coordination trials). Based on the high accuracy, all trials were analyzed for the EEG analysis.

The mean (± SD) reaction time of correct responses was 1617 ± 345 ms when the agent was pinged, 1616 ± 357 ms when the patient was pinged, 1617 ± 345 ms for all event trials and 1149 ± 349 ms for all coordination trials. The reaction times reported here were first averaged within subjects for all the correct trials, then averaged across subjects. Paired t-test (two-tailed) indicated no significant difference between the reaction times for agent ping trials and patient ping trials, *t*(15)=0.6, *p*=0.99. Paired t-test (two-tailed) revealed a significant difference between the reaction times for event trials and coordination trials, *t*(15)=9.7, *p*<0.001, similar to the results in Experiment 1.

#### 2.2.2. EEG results

Similar to Experiment 1, a cluster-based permutation test (Maris & Oostenveld, 2007) was performed on the responses to the event picture and the coordination picture in the time window of -200:3500 ms for the NSW ROI using customized MATLAB scripts. A cluster was identified in the negative slow wave (NSW) ROI, extending from 637 to 1109 ms after picture onset (*p*=0.006, Figure 8). As in Experiment 1, the difference manifested as an increased negativity for the event pictures relative to the coordination pictures in these posterior electrodes. The scalp topography of this NSW effect is generally similar to the one obtained in Experiment 1. The duration of the sustained effect appeared to extend just up through the onset of the ping screen, after which no significant differences were observed.

**Figure 8.**
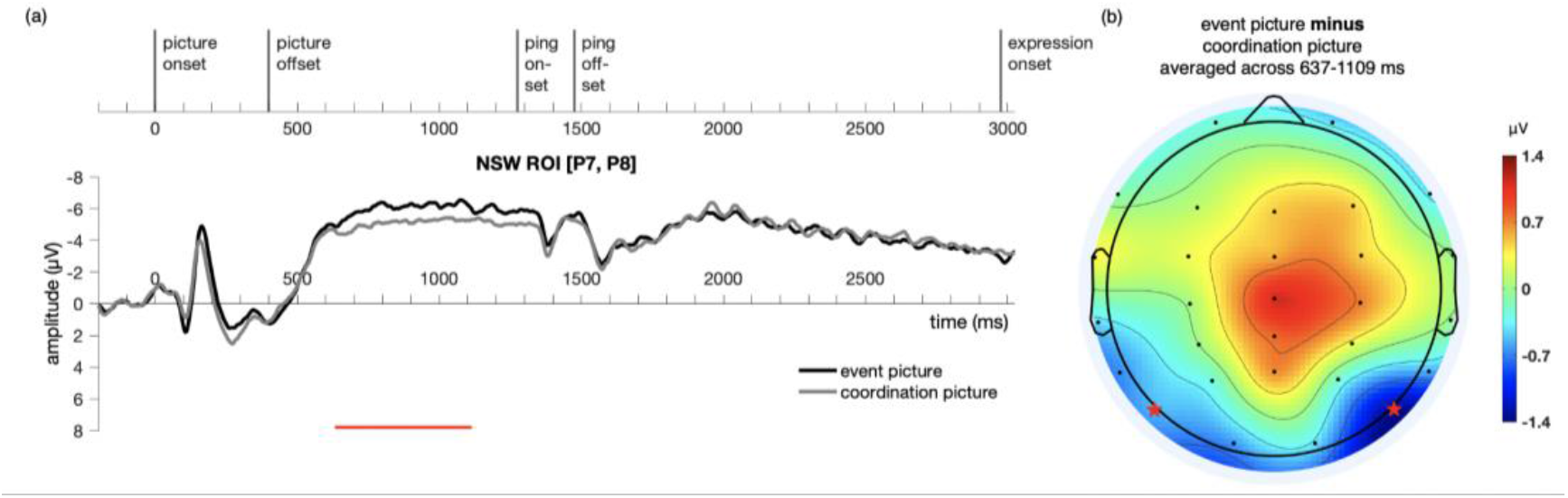
Illustration of the results for epochs time-locked to picture onset in the NSW ROI (electrodes P7 and P8, as marked by the stars in Figure b) for Experiment 2. (a) The significant cluster (637-1109 ms) is marked in red. (b) The scalp distribution average across the 637-1109 ms time window. NSW: negative slow wave; ROI: region of interest.

ERP responses to the delay-period agent ping and the patient ping in the event condition are illustrated in Figure 9. Similar to Experiment 1, cluster-based permutation test (Maris & Oostenveld, 2007) was performed comparing the agent ping vs. patient ping conditions in the time window of -200:1800 ms using customized MATLAB scripts using the same approach described above for the event picture comparison. No significant clusters distinguishing the agent ping and patient ping were identified in the CPP ROI, nor in the NSW ROI.

**Figure 9.**
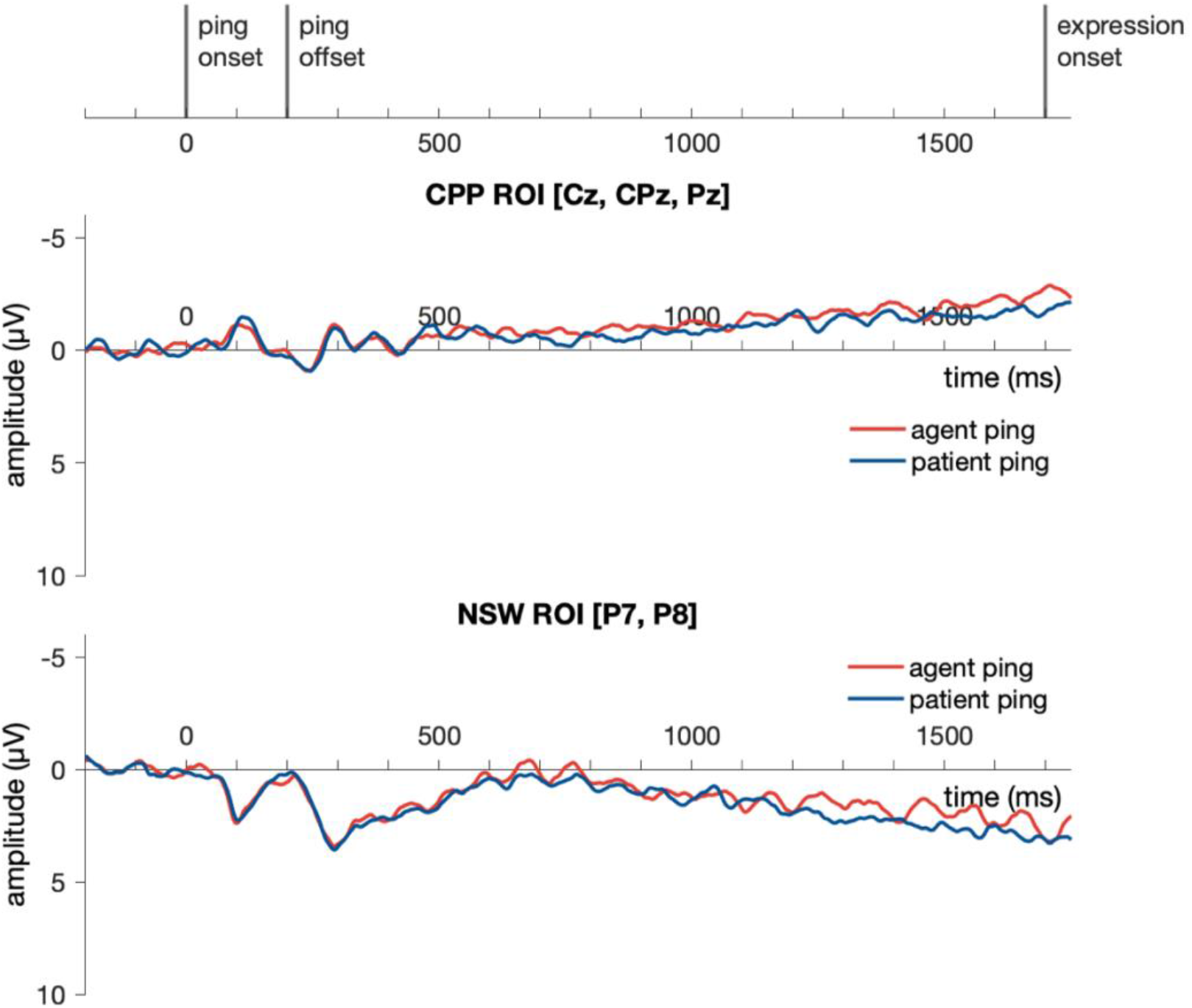
Illustration of the results for epochs time-locked to ping onset (−200:1800 ms) in the NSW ROI and the CPP ROI for Experiment 2. NSW: negative slow wave; CPP: centro-parietal positivity; ROI: region of interest.

## 3. Discussion

Our current experiments provide one of the first electrophysiological studies directly addressing how event relations and event roles (i.e. agent and patient) are maintained in working memory. We report two key findings. First, across Experiments 1 and 2, we saw that the working memory retention of events (e.g., a tiger hit an elephant) relative to simple coordination (e.g., a tiger and an elephant) was hallmarked by a response similar to a negative slow wave (NSW) in posterior-occipital electrodes during the delay period. Events were more negative compared to coordinations, suggesting higher working memory load maintaining events than coordinations, based on standard interpretations of the negative slow wave (for a review see Section 1.1.2). Experiment 2 confirmed this sustained difference from the onset of the delay period. Our second finding was that in a novel agent vs. patient “pinging” manipulation during the delay, we observed no reliable difference over centro-parietal positivity electrodes or negative slow wave electrodes for agent as compared to patient. Interestingly, however, the sustained negative slow wave effect seemed to be largely dampened after the presentation of the ping, as observed most clearly by means of the shift of the ping presentation timing in Experiment 2.

### 3.1. Representing event relations in working memory

The major difference between the event and coordination conditions is whether or not an event relation needs to be represented across the delay period. The NSW effect in Experiment 1 onset at ∼ 450 ms and extended even until ∼ 2500 ms; for Experiment 2, it onset at ∼ 600 ms and extended until ∼ 1100 ms (approximately when it was interrupted by the ping). This late, *sustained* effect across the two conditions suggests that event relations *can* be represented by sustained activity in working memory (crucially, working memory representation does not need to be supported by sustained activity, see Section 3.3 below). Our current study marks a crucial first step towards further elucidating the implementation-level question of *how* event relations are represented in working memory.

Where in the brain may be generating the NSW effects in our current study? Being a late sustained effect, these time windows are comparable to the trend in the posterior parietal cortex in Hultén et al. (2014; onset at around 300 ms and extends until at least 1000 ms). Therefore, we speculate that the NSW effect in our current experiments reflects sustained activity in the posterior parietal cortex; this is consistent with previous reports on the contribution of the posterior parietal cortex in working memory (Robitaille et al., 2010; Z. Fan et al., 2012; Becke et al., 2015; Bettencourt & Xu, 2016; K. Jia et al., 2021) and event representation (Thompson et al., 2007; Centelles et al., 2011; J. Wang et al., 2016; Williams, Reddigari & Pylkkänen, 2017; Matchin et al., 2019a, b). Moreover, the posterior parietal cortex has been observed to be involved in both linguistic and visual processing, suggesting that it is hosting conceptual representations (Jouen et al., 2015; Zadbood et al., 2017; Baldassano et al., 2017; Menenti, Segaert & Hagoort, 2012; Liu et al., 2020; Ivanova et al., 2021). Taken together, we believe that the NSW effect observed here is likely reflecting object indexicals at the conceptual level.

Previous work (Summerfield, Luyckx & Sheahan, 2020) has proposed that the posterior parietal cortex is representing “relational structures” (see also O’Reilly, Ranganath & Russin, 2022), yet the format of this representation remains elusive. One possibility consistent with the negative slow wave (NSW) effect during the maintenance of events vs. coordinations is that the event relation in an event is represented by a separate object indexical apart from the two participants. These “relation indexicals” could be qualitatively different from the indexicals for objects; a possibility that could be tested in future experiments. However, we note that it is also possible that the NSW we observed was driven by a non-indexical representation of event relations by other regions in the posterior parietal cortex (or even other brain regions). Further work is needed to disentangle these possibilities.

Interestingly, Lapinskaya et al. (2016) observed a late (from 500 ms to at least 800 ms) parietal-occipital ERP effect between object nouns vs. event nouns and verbs after the presentation of the second word in a two-word similarity judgement task. This effect had a similar scalp distribution to our current effect based on visual inspection. The object nouns were more positive over the posterior electrodes compared to verbs and event nouns, which mirrors the current results in which coordinations elicited a less negative response than events. However, Lapinskaya et al. (2016) used a single-word similarity judgment paradigm, which is quite different than the paradigm of our current study, and did not examine responses over an extended maintenance delay.

We should also note that some behavioral studies have shown that when two objects are involved in an event, participants can actually perform better on working memory tasks than when they are maintaining two separate objects (Stahl & Feigenson, 2014; Ding, Gao & Shen, 2017; Paparella & Papeo, 2022; Lu et al., 2022). This seeming contradiction with our current results remains to be resolved in future research. One possibility is that the number of indexicals consumed in working memory for the same input may vary according to task environments, as suggested by studies on visual grouping (Balaban & Luria, 2016; Rabbitt et al., 2017; McCants, Katus & Eimer, 2020). Another possibility is that different numbers of indexicals may be consumed during extraction and maintenance phases, as suggested for visual grouping (Peterson et al., 2015; Thyer et al., 2022).

### 3.2. Representing event roles in working memory

In this study we introduced a novel “pinging” paradigm to attempt to observe evidence of the differential coding of event roles in working memory across the delay. One hypothesis was that event roles are coded similarly to magnitudes, and that this would be associated with differences in the electrodes previously associated with a centro-parietal positivity electrodes suggested to reflect magnitude coding. We failed to find evidence for this hypothesis, as we did not observe a significant effect of the pinged event role over CPP electrodes across the two experiments. This is not definitive evidence against magnitude coding of event roles, as the CPP may not be a general hallmark for all magnitudes. We also note that much of the existing pinging literature has relied on decoding but not ERP comparison. Notably, Fornaciai & Park (2020), in a numerosity working memory task, identified a significant ERP difference upon sample presentation across conditions, yet failed to identify significant ERP differences upon ping presentation during the retention period across these conditions. In line with the rest of the pinging literature, they observed above-chance decodability across conditions upon ping presentation. The potential absence of ERP difference upon ping presentation in our current study is in line with Fornaciai & Park (2020). Further research is needed to investigate whether ERPs are generally insensitive to the neural responses modulated by pinging paradigms.

On the other hand, there are also theoretical challenges for representing event roles as magnitudes. For example, if we consider tools or instruments (e.g., Mary opened the bottle with *the opener*), it is hard to think of this role as being somewhere on a 1-D scale along the same line of agent and patient. This is different from other magnitudes (e.g., numerosity, magnitude of reward, and magnitude of evidence during decision-making), which can be understood as being represented on a 1-D magnitude scale. What are the alternative mechanisms beyond magnitude underlying event role representation? There are currently at least two other major hypotheses about the implementation of event roles. One is that different event roles, as features, are represented by different neural firing patterns. The other is that agent and patient roles are signaled by the most excited phase of this object indexical in an oscillation. For example, when an object indexical is representing the agent object, it is most excited at a certain phase, while when the same object indexical is representing the patient object, it is most excited at another phase. The brain may make use of both representations for different purposes. The first hypothesis is supported by the fMRI decoding study of J. Wang et al. (2016), which successfully decoded above chance whether a certain animal is the agent or the patient in a visual scene from a wide range of brain regions. Some forms of the second hypothesis have been discussed in the computational literature for a long time (Shastri & Ajjanagadde, 1993; Shastri, 1999, 2002, 2007; Hummel & Holyoak, 2003; A. E. Martin & Doumas, 2017), yet have only recently been tested by Peng et al. (2017). Peng and colleagues, using the dense sampling method for identifying behavioral oscillations, found that the object indexicals for the agent objects and the patient objects are most excited at a ∼ 90 degree lag in the alpha band (∼ 10 Hz). Further studies need to be conducted in order to draw further connections between the object indexical system and event role representation from an oscillation point of view. Specifically, studies up to date have been in line with the possibility that when holding multiple objects (≥ 2) in working memory, different object indexicals are most excited at different phases of an oscillation (J. Jia et al., 2017; Peters et al., 2021; Pomper & Ansorge, 2021; Balestrieri, Ronconi & Melcher, 2021; Chota et al., 2022). This leaves open the possibility that the event role of an object adjusts the most excited phase of its corresponding object indexical, with different event roles adjusting the phase in different manners; this remains to be tested in future research. Our current results do not provide strong evidence either for or against these hypotheses.

Another dispute over the representation of event roles is whether they are independent or dependent from the event type (see Williams, 2015, 2021). By relation-independent, we refer to the abstract roles like agent and patient; by relation-dependent, we refer to the action-specific roles such as hitter and hittee, hugger and huggee, etc. For relation-dependent roles, there are also two possibilities: one is that different types of agents are aligned on a dimension of “agentivity” (for example, hitters may be more “agenty” than huggers), and the other is that there is no such dimension of “agentness” or “patientness”, i.e. there is no systematic change in the representational patterns for more “agenty” agents or more “patienty” patients. While a large body of literature in formal semantics assumes the existence of relation-independent roles (for discussions see Williams, 2021), Dowty (1991) laid out some features that are typical for (proto-)agents and (proto-)patients respectively, which can be empirically quantified by subjective ratings that are graded in nature (Kako, 2006a, b; Reisinger et al., 2015; Rissman & Lupyan, 2022, for studies with a similar logic using other ratings see Johnson, 1967; Madnani, Boyd-Graber & Resnik, 2010). Indeed, it is possible to have both relation-dependent and relation-independent representations of event roles in the brain (Frankland & Greene, 2020), yet more research is still needed to resolve this dispute.

### 3.3. Sustained activity vs. activity-silent mechanisms

Our current results are consistent with the idea that event relations can be maintained in working memory in the form of sustained neural activity, given that we observe a sustained ERP difference between the retention of events and coordinations. Although sustained activity has been seen as a key mechanism for working memory representation for a long time (for reviews see Leavitt, Mendoza-Halliday & Martinez-Trujillo, 2017; X. J. Wang, 2021), at least two lines of research have argued that working memory retention can also sometimes be implemented by “activity-silent mechanisms” (Lewis-Peacock et al., 2012; Stokes, 2015; Rose et al., 2016; Kamiński & Rutishauser, 2020; Beukers et al., 2021) or other mechanisms that do not induce sustained changes in neural metabolism (e.g., Wutz, Zazio & Weisz, 2020; Lundqvist et al., 2021).

One line of research has utilized visual stimuli, and observed that the amplitude of CDA after presenting two consecutive memory arrays is not always reflective of the load of the first memory array (Berggren & Eimer, 2016; Feldmann-Wüstefeld, Vogel & Awh, 2018; J. Zhang et al., 2021) and that a huge disruption in sustained activities does not necessarily accompany a huge impairment in memory performance (Kreither, Papaioannou & Luck, 2022). Another line of research has utilized linguistic stimuli, and observed that the maintenance of syntactic dependency does not always guarantee the presence of the sustained component of SAN (sustained anterior negativity) (McKinnon & Osterhout, 1996; Kaan et al., 2000; Lau & Liao, 2018; Yano & Koizumi, 2018, 2021; Cruz Heredia, Dickerson & Lau, 2021; Sprouse et al., in preparation; Lo & Brennan, 2021; Lau & X. Yang, 2023; for an earlier review see Lau, 2018). This line of work implies that syntactic dependencies rely on a form of working memory that does not require sustained neural activity.

Our current experiment provides one of the first reports about how event relations are represented across time in working memory. The sustained negative slow wave effect across events and coordinations in Experiment 1 and 2 demonstrated that the working memory maintenance of event relations *can* be realized by sustained activity. However, as Experiment 2 confirmed, this sustained NSW effect was largely dampened by the ping. This is in line with previous findings in visual working memory studies that an interruption during retention dampened CDA too (Berggren & Eimer, 2016; Feldmann-Wüstefeld, Vogel & Awh, 2018; J. Zhang et al., 2021), yet resulting in little detriment to behavioral accuracy (Kreither, Papaioannou & Luck, 2022)^6^. This suggests that the representational format of the memorandum shifted from sustained activity to activity silent representations upon the interruption, which may also be the case for our current experiments. Therefore, our observation that the sustained difference was reduced after the ping suggests that although the working memory maintenance of event relations *can* be realized by sustained activity, it does not have to and can be represented in an activity silent format.

### 3.4. Event type and social interactions

In the current study, we focused on how the brain reacts to whether or not there is an event relation at all in working memory, but not how the event type (e.g., hitting, kicking) is represented. Previous fMRI studies have converged on the posterior lateral temporal cortex (PLTC, Wurm & Lingnau, 2015; Wurm et al., 2016; Hafri, Trueswell & Epstein, 2017; Walbrin & Koldewyn, 2019; Wurm & Caramazza, 2019) as hosting the representation for event types. Following our interpretation of the negative slow wave effect, it is possible that the indexical for event relation in the PPC (posterior parietal cortex) points to the event type represented in PLTC during working memory maintenance. However, evidence remains mixed on whether the representations in PLTC about event type reflect the extraction of event types or the maintenance of event types in working memory, or both (Lu et al., 2016; H. Zhou et al., 2022).

The NSW effect in our current study can also be interpreted as reflecting social cognition computations in particular, related to the presence of social interaction for events. Future research remains to be conducted in order to elucidate whether all event types can drive an NSW effect, or only social ones can. As we reviewed in the previous paragraph, PLTC hosts the representation for event types. Interestingly, some recent studies have suggested that some sub-regions in PLTC may favor social interactions more than non-social ones (Wurm, Caramazza & Lingnau, 2017; Wurm & Caramazza, 2019; H. Yang et al., 2020).

### 3.5. Conclusions

The current study provides one of the first attempts to discover ERP hallmarks for the conceptual representation of events vs. coordination and agent vs. patient roles in working memory. We observed a sustained difference during a delay period for maintaining conceptual information about events relative to coordinations in working memory, which resembled the posterior-occipital negative slow wave (NSW) effect observed in previous visual working memory studies. Interestingly, a novel event role “pinging” manipulation also revealed that the NSW effect seemed to be largely dampened after the presentation of the ping.

Much future work still needs to be done to gain a better understanding of these electrophysiological hallmarks observed in our current study, in order to better understand the format of event representation in working memory. Our current work is an exploratory yet necessary first step.

## 4. Materials and methods

### 4.1. Experiment 1

#### 4.1.1. Participants

In total, 24 subjects participated in this experiment; they were balanced in terms of the list used in the main experiment. Of these subjects, data from 5 subjects were excluded: 1 of these 5 subjects was excluded because of incomplete data. Data from the other 4 of the 5 subjects were excluded based on a procedure detailed in Section 4.1.5. The remaining 19 subjects analyzed had an age of 18-27 (*M*=22, 11 female), all right-handed as measured by a translated version of the Edinburgh Handedness Inventory (Oldfield, 1971). Subjects were recruited from the New York University Shanghai and East China Normal University community. Subjects received monetary reimbursement for their participation. Written consent was acquired from each subject, and the procedures were approved by the Institutional Review Board of the University of Maryland and New York University Shanghai.

#### 4.1.2. Stimuli

144 “event pictures” about events involving two characters were constructed, adapted from stimuli used in Hultén et al. (2014). The Hultén et al. (2014) stimuli originally consisted of cartoons depicting a single animal conducting a certain action or standing still (side-facing, used as the patients in event pictures, or front-facing, used in coordination pictures; for the definition of coordination pictures see below).

The characters used in the stimuli set were four animals (the elephant 大象, the lion 狮子, the hippo 河马 and the sheep 绵羊, therefore, there are 6 combinations). Each event involved one of 6 distinct actions (hitting 打, pulling 拉拽, kissing 亲, pointing at 指, pushing 推, scolding 骂). From each of the 6 character duos (e.g., the elephant and the lion), each of the 6 actions (e.g., hitting), and each of the 2 ways of allocating event roles (e.g., the elephant hitting the lion vs. the lion hitting the elephant), we constructed 72 unique event pictures. Note that due to the nature of the Hultén et al. (2014) stimuli (the animals performing an action were all right-facing), the agent was always to the left in these 72 “original” pictures. Therefore, we created another 72 mirror-flipped pictures by mirror-flipping the 72 original pictures to balance the relative visual position of the two characters, thus resulting in 144 event pictures in total.

Another 12 “coordination pictures” were also constructed, with two characters (also within the elephant, the lion, the hippo and the crocodile) standing next to each other. For each animal pair (X,Y), we created separate pictures in which animal X is to the left or animal Y is to the left.

The same event (e.g., lion hitting elephant) was only presented once for each subject. In order to balance relative right/left positions, we distributed the 144 original and mirror-flipped pictures across two lists (list A and B), each containing 72 pictures representing unique events; in each list, a certain character had an equal probability of appearing to the left as agent, to the left as patient, to the right as agent, and to the right as patient. Besides, for each action, the probability of the agent being to the left was the same as the probability of the agent being to the right. Both list A and list B contained an equal amount of original (i.e., 36) and mirror-flipped (i.e., 36) event pictures. Each list also included 72 coordination pictures (6 repetitions of each of the 12 coordination pictures). That is, each of the two lists consisted of 144 pictures in total (72 event pictures and 72 coordination pictures); this corresponds to 144 trials that each subject went through (72 event trials and 72 coordination trials). For more examples of our picture stimuli, see Figure S2.

Each trial contained a single word “ping” stimulus during the delay between the picture and the probe linguistic expression. These stimuli were arranged so that within each list, (a) there was an equal probability for the agent on the left, the agent on the right, the patient on the left and the patient on the right to be pinged, (b) for each action, the agent and patient were equally likely to be pinged, (c) for each pinged character, there was an equal possibility that this character was the agent or the patient.

The format of probe linguistic expressions is illustrated in Table 1. In trials where an event picture (e.g., a picture depicting a lion hitting an elephant) had been presented (i.e. event trials), the linguistic expression (here, a sentence) either matched with the event picture (50% probability, e.g., “the lion hit the elephant”/”the elephant was hit by the lion”) or was different in event role assignment compared to the event picture (50% probability, e.g., “the elephant hits the lion”/”the lion was hit by the elephant”). The sentence was determined randomly to be in active or passive form. In trials where a coordination picture (e.g., a picture depicting a lion standing next to an elephant) had been presented (i.e. coordination trials), this linguistic expression (here, a phrase) either matched with the picture (50% probability, e.g., “the lion and the elephant”/”the elephant and the lion”) or was different in character compared to the coordination picture (50% probability, e.g., “the lion and the hippo”, “the elephant and the hippo”, etc.). The order of the two animals was determined randomly.

**Table 1.**
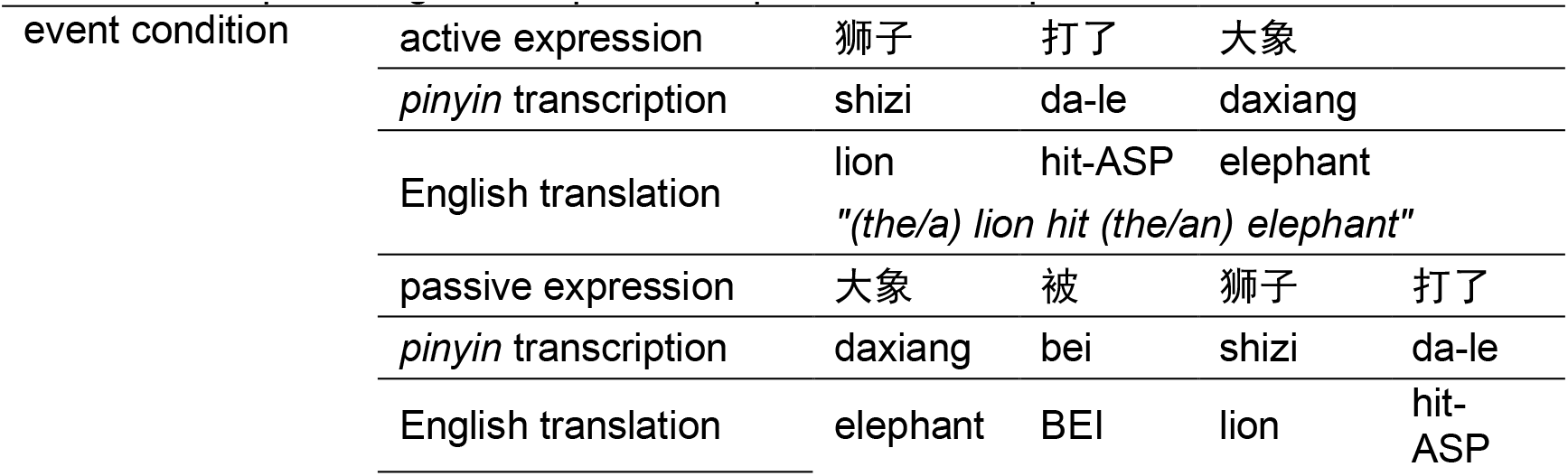
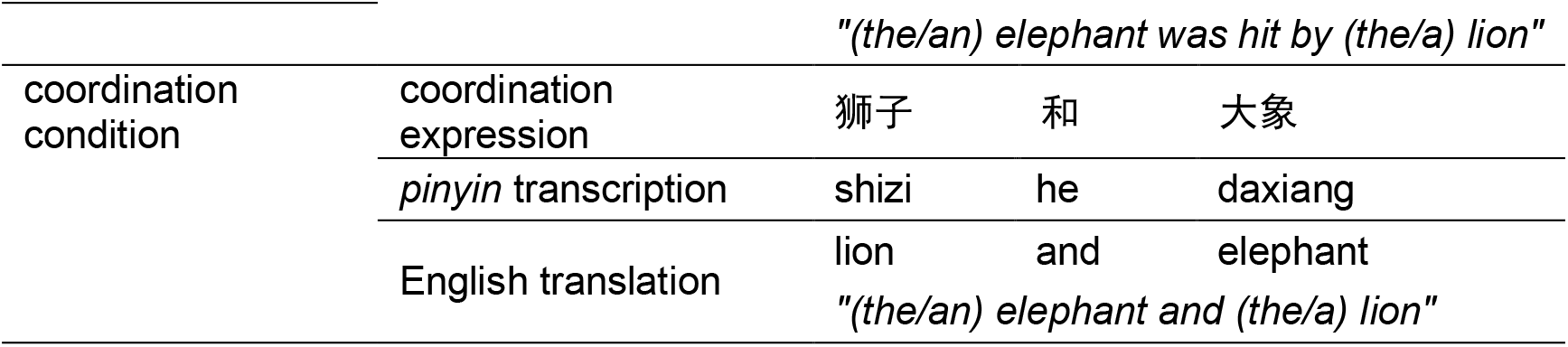
Example of linguistic expressions presented in Experiment 1.

#### 4.1.3. Procedure

Each trial started with a “+” in the middle of the screen (for 500 ms) signaling the beginning of a trial. This “+” was followed by the presentation of an event picture for 1000 ms (for example, a picture depicting the lion hitting the elephant) or a coordination picture (for example, a picture depicting the lion standing next to the elephant). After 900 ms of a blank screen, a brief “pinging” word in larger font (250 pt) referring to one of the two characters (e.g., “lion” or “elephant”) was presented for 200 ms, in order to highlight the just-encoded character-role pairing maintained in working memory. This ping was followed by a blank screen for 600 ms, resulting in a total SOA (stimulus onset asynchrony) of 800 ms between ping presentation and the subsequent probe linguistic expression (a sentence or a phrase). After the delay period, a linguistic expression (see Table 1) appeared at the center of the screen, which either matched or did not match with the picture. The whole linguistic expression was presented on the same screen.

Subjects were asked to make a response on the keyboard whether the linguistic expression matched with the picture in the same trial as fast and accurately as possible; subjects were allocated into two response hand groups: in one group subjects pressed “z” with the left hand when matching and pressed “m” with the right hand when not; in the other group subjects pressed “m” with the right hand when matching and pressed “z” with the left hand when not. Then a feedback screen displaying whether the choice was right (in green) or wrong (in red) was presented for 400 ms; if the subjects did not make the response within 5 s, the feedback screen would automatically appear, showing “no response” (in white). After the feedback screen, there was an inter-trial interval (ITI) uniformly random across 1000-2000 ms before the next trial. Within the ITI, a line of asterisks appeared on the screen; before the experiment, the subjects were encouraged to blink during this period, in order to reduce blinks within trials. Subjects were instructed to always fixate on the center of the screen within a trial. For a schematic illustration of one trial in Experiment 1 see Figure 10.

**Figure 10.**
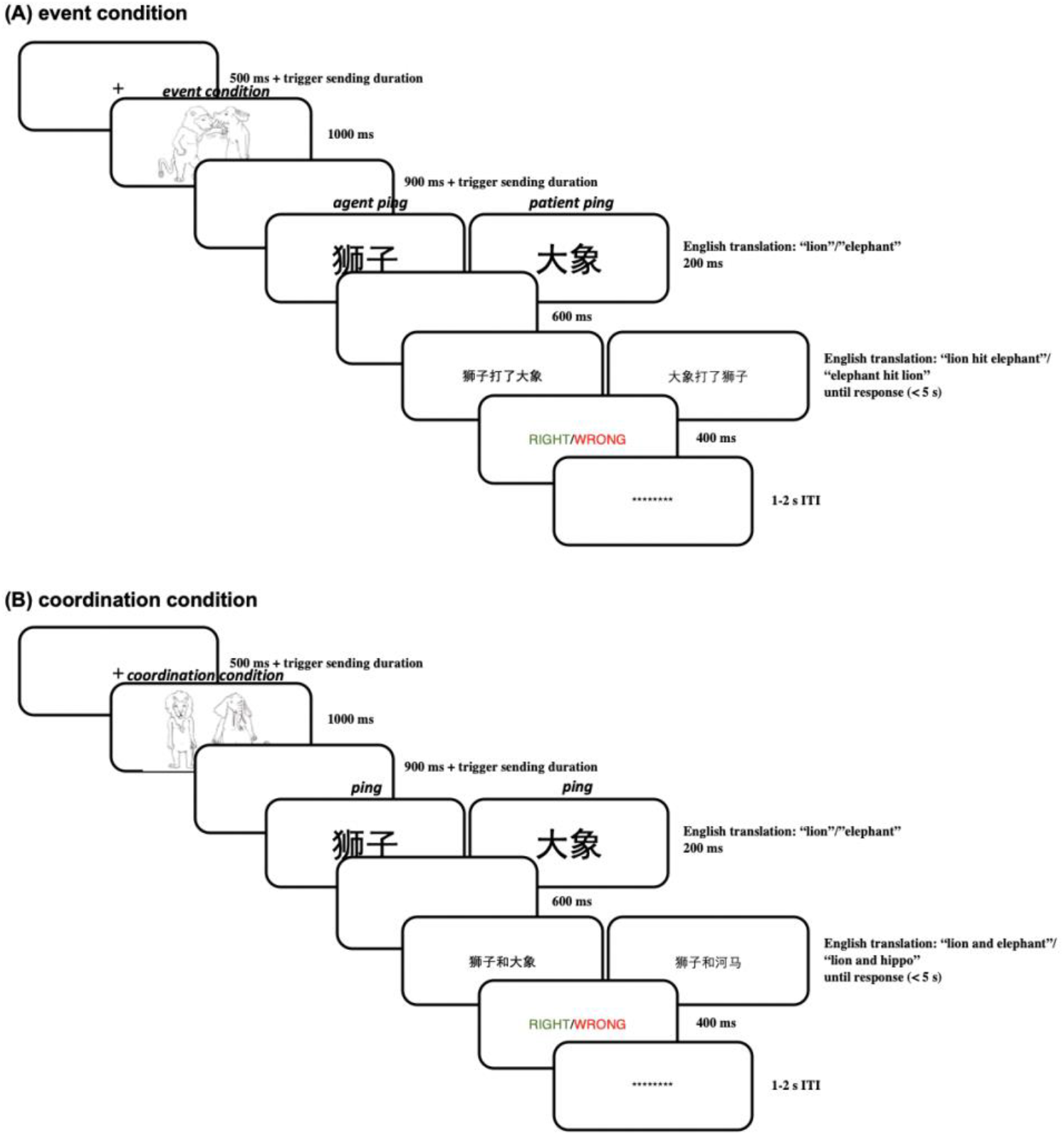
(A) Illustration of one trial in the event condition in Experiment 1; (B) illustration of one trial in the coordination condition in Experiment 1.

**Figure 11.**
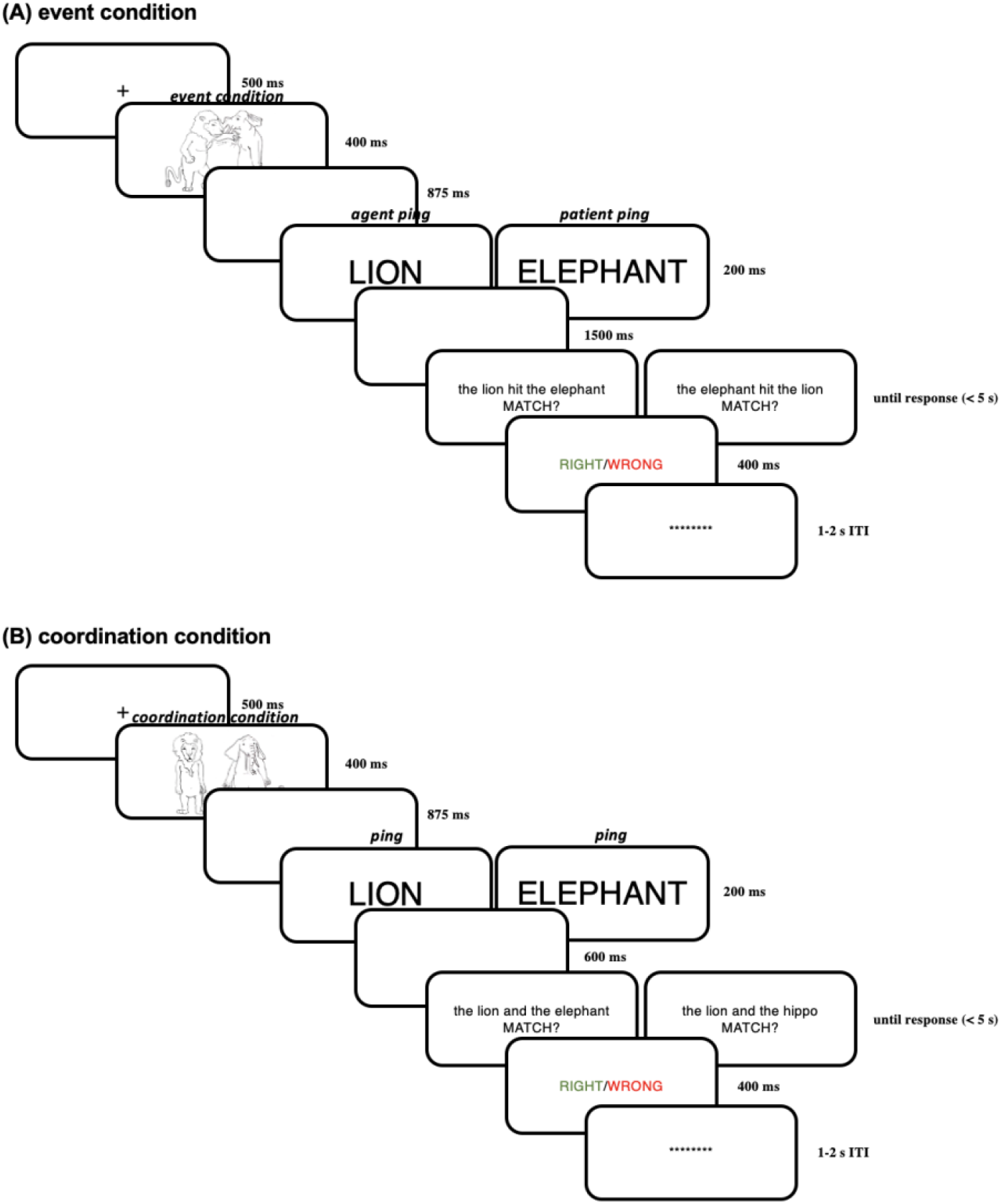
(A) Illustration of one trial in the event condition in Experiment 2; (B) illustration of one trial in the coordination condition in Experiment 2.

Before the main experiment, subjects were given 24 practice trials (12 event pictures, 12 coordination pictures) randomly, constructed from two animals not involved in the experiment (which is, the cow 奶牛 and the crocodile 鳄鱼) and all 6 actions. Two balanced lists of event pictures (each containing 12 event pictures) were similarly created. The practice trials were used to familiarize subjects with the verbs used for the actions, as well as the response for coordination sentences, etc. These 24 practice trials were administered repetitively until the subjects reached 100% accuracy on all 24 practice trials. Note that in practice trials, the unmatching linguistic expressions for coordination pictures were constructed by substituting one of the characters (i.e. the cow or the crocodile) for another animal that was not involved in this experiment (which is, the squirrel 松鼠). This difference in the probe phrases compare to our main experiment was unlikely to affect our results, as this does not change essentially that in the coordination conditions, subjects were representing the two characters as two grouped entities.

In the main experiment, five self-paced breaks were given every 24 trials (there were in total 144 trials). The experiment was run with Psychtoolbox 3 (Brainard, 1997; Kleiner, Brainard & Pelli, 2007) in MATLAB 2020b. The background of the screen was grey, and the texts were white. The pictures were created in 1600 × 1200 px, and were presented on the screen as 24° × 18°. All texts, except for the ping, were presented with a font size of 100 pt; the pings were presented with a font size of 250 pt. Subjects sat at a distance of 80 cm before the screen.

#### 4.1.4. Behavioral data analysis

The reaction times were first averaged within subjects for all the correct trials, then paired t-tests were conducted across conditions in JASP 0.16.1 (JASP Team, 2022).

#### 4.1.5. EEG recording

EEG signal was recorded using a 32-channel active electrode system (Brain Vision actiCHamp; Brain Products) with a 1000 Hz sampling rate in an electromagnetically shielded and sound-proof room. Electrodes were placed on an EasyCap, on which electrode holders were arranged according to the 10-20 international electrode system. The impedance of each electrode was kept below 10 kΩ. The data were referenced online to electrode Cz. Two additional EOG electrodes (HEOG and VEOG) were attached for monitoring ocular activity. The EEG data were acquired with Brain Vision PyCoder software (http://www.brainvision.com/pycorder.html) and filtered online between DC and 200 Hz with a notch filter at 50 Hz.

#### 4.1.6. EEG data analysis

EEG data were analyzed using customized MATLAB (version: R2020b) scripts based on EEGLAB v2021.0 (Delorme & Makeig, 2004) and ERPLAB v8.20 (Lopez-Calderon & Luck, 2014). Bad channels (identified by visual inspection, 0-1 channels for each subject) were first interpolated (spherical); throughout the whole analyzing pipeline, only scalp electrodes were included. Then the EEG data were re-referenced using average (because the ICLabel plug-in was trained on average-referenced data, Pion-Tonachini, Kreutz-Delgado & Makeig, 2019), then down-sampled to 256 Hz, using functions in EEGLAB. Then the data were filtered by an IIR Butterworth filter with a cut-off at 0.01 Hz and 40 Hz using functions in ERPLAB (DC bias was also removed). Then ICA was conducted with the runica algorithm, and components that were labelled (using the ICLabel plug-in, Pion-Tonachini, Kreutz-Delgado & Makeig, 2019) with labels “Muscle”, “Eye”, “Heart”, “Line Noise”, or “Channel Noise” with a confidence of ≥ 90% were removed from data. Each subject had 0 to 2 components removed.

Epochs of -500:3500 ms were extracted (time locked to picture onset or ping onset), baseline corrected using the period -200 ms to 0 before picture onset. Epochs in which at least one electrode had a range of more than 100μV during the time interval of interest (picture: -200:3500 ms, ping: -200:1800 ms) were excluded. For each subject, if there was at least one condition (i.e. event picture, coordination picture, agent ping, patient ping) that had fewer than 20 epochs, this subject was excluded. This criterion excluded the 4 subjects mentioned in Section 4.1, resulting in 19 subjects analyzed.

For the event picture vs. coordination picture comparison, we selected only - 200:3500 ms epochs time-locked to picture onset that passed the above exclusion criteria (a range of no more than 100μV) in these 19 subjects. We first averaged the ERPs within subjects across epochs for each electrode. For the NSW ROI analysis, we averaged the response in the NSW ROI (P7 and P8) for each subject, then conducted cluster-based permutation test across time (Maris & Oostenveld, 2007) by the permutest() function (Gerber, 2021) comparing the response for the NSW ROI for event picture vs. coordination picture conditions. In the cluster-based permutation tests, first, potential clusters were identified with point-to-point paired t-test at a threshold of alpha = 0.05 (two-tailed). Then, the t values of these potential clusters were compared against a permuted (permutations = 9999) distribution of t values, with a threshold of alpha = 0.05 (two-tailed).

For the agent ping vs. patient ping comparison, we selected only -200:1800 ms epochs time-locked to ping onset that passed the above exclusion criteria (a range of no more than 100μV) in these 19 subjects. We first averaged the ERPs within subjects across epochs for each electrode. For the CPP ROI analysis, we averaged the response in the CPP ROI (CP1, CP2, Pz) for each subject, then conducted cluster-based permutation test (Maris & Oostenveld, 2007) by the permutest() function (Gerber, 2021) comparing the response for the CPP ROI for agent ping vs. patient ping conditions. Because we did not observe significant clusters in this pre-determined analysis, we then exploratorily conducted a cluster-based permutation test across time with the same parameters in each of the electrodes separately.

After beginning data analysis, we discovered an inaccuracy in the timing parameters of Experiment 1 which introduced jitter between the actual presentation of the picture and the recording of the event by the EEG system. Fortunately, a photodetector signal accompanied the ping presentation, allowing a precise estimation of its actual timing. We used this validated time estimate for the ping presentation to infer the timing of the picture presentation, but this means the estimate of the picture onset timing is less direct. In the Experiment 2 replication we used a different presentation code and oscilloscope testing to ensure precise temporal alignment of the trigger codes for both picture presentation and ping presentation.

### 4.2. Experiment 2

#### 4.2.1. Participants

In total, 23 subjects participated in this experiment. Of these subjects, data from 7 subjects were excluded: 2 subjects was excluded because of incomplete data; 1 subject was excluded because of low behavioral accuracy (63%); 4 subjects were excluded based on the same procedure as Experiment 1, detailed in Section 4.1.5. The remaining 16 subjects had an age of 18-30 (*M*=24, 8 female). 13 of them were right-handed and 3 of them were left-handed, as measured by the Edinburgh Handedness Inventory (Oldfield, 1971). They were balanced in terms of the list used in the main experiment, the list used in the practice session, and response hand. Subjects were recruited from the University of Maryland, College Park community. Subjects received monetary reimbursement for their participation. Written consent was acquired from each subject, and the procedures were approved by the Institutional Review Board of the University of Maryland.

#### 4.2.2. Stimuli

The picture stimuli in Experiment 2 were the same as Experiment 1. The probe linguistic expressions in Experiment 2 were the same, except that they were in English: e.g., “the lion pointed at the elephant”, “the sheep was hit by the lion”, “the lion and the hippo”. The pings in Experiment 2 were English words in capital letters, e.g., “LION”.

#### 4.2.3. Procedure

The procedure of Experiment 2 is generally similar to Experiment 1, except for the timings and font sizes (modified because of a smaller screen). Each trial started with a “+” in the middle of the screen (for 500 ms) signaling the beginning of a trial. This “+” was followed by the presentation of an event picture for 400 ms or a coordination picture. After 875 ms of a blank screen, a brief “pinging” word in capital letters and larger font (100 pt) referring to one of the two characters (e.g., “LION” or “ELEPHANT”) was presented for 200 ms, in order to highlight the just-encoded character-role pairing maintained in working memory. This ping was followed by a blank screen for 1500 ms. After the delay period, a linguistic expression in English appeared at the center of the screen, which either matched or did not match with the picture. The whole linguistic expression was presented on the same screen.

Subjects were asked to make a response on the keyboard whether the linguistic expression matched with the picture in the same trial as fast and accurately as possible. Then a feedback screen displaying whether the choice was right (in green) or wrong (in red) was presented for 400 ms; if the subjects did not make the response within 5 s, the feedback screen would automatically appear, showing “no response” (in white). After the feedback screen, there was an inter-trial interval (ITI) uniformly random across 1000-2000 ms before the next trial. Within the ITI, a line of asterisks appeared on the screen; before the experiment, the subjects were encouraged to blink during this period, in order to reduce blinks within trials. Subjects were instructed to always fixate on the center of the screen within a trial. For a schematic illustration of one trial in Experiment 2 see Figure 10.

The background of the screen was grey, and the texts were white. The pictures were presented on the screen as 24° × 18°. All texts, except for the ping, were presented with a font size of 30 pt; the pings were presented with a font size of 100 pt. The distance between the subject and the screen was measured individually.

#### 4.2.4. Behavioral data analysis

Same as Experiment 1.

#### 4.2.5. EEG recording

Twenty-nine tin electrodes were held in place on the scalp by an elastic cap (Electro-Cap International, Inc., Eaton, OH) in a 10-20 configuration. Bipolar electrodes were placed above and below the left eye and at the outer canthus of both eyes to monitor vertical and horizontal eye movements. Additional electrodes were placed over the left and right mastoids. Scalp electrodes were referenced online to the left mastoid. The ground electrode was positioned on the scalp in front of Fz. Impedances were maintained at less than 20 kΩ for all scalp electrode sites, and less than 10 kΩ for mastoid and ocular electrodes. The EEG signal was amplified by a NeuroScan SynAmps® Model 5083 (NeuroScan, Inc., Charlotte, NC) with a bandpass of 0.05–100 Hz and was continuously sampled at 500 Hz by an analog-to-digital converter.

#### 4.2.6. EEG data analysis

The analysis pipeline was the same as Experiment 1 (Section 4.1.6), except that for Experiment 2, the CPP ROI was Cz, CPz and Pz. There were 0-3 interpolated bad channels for each subject, and 0-1 ICA component was removed for each subject.

## Acknowledgements

This work is supported by NSF #1749407 (to E.L.). We would like to thank Matti Laine for sharing the stimuli from Hultén et al. (2014).

## Author contributions

Xinchi Yu: Conceptualization, data curation, investigation, formal analysis, methodology, software, visualization, writing-original draft, writing-review & editing

Jialu Li: Data curation, investigation, software, writing-review & editing

Hao Zhu: Data curation, investigation, software, writing-review & editing

Xing Tian: Resources, supervision, writing-review & editing

Ellen Lau: Conceptualization, funding acquisition, methodology, supervision, writing-original draft, writing-review & editing

## Footnotes

1 Note that there are many types of features. For example, one can make the distinction between perceptual (e.g., color, shape) and conceptual (e.g., occupation, edibility) features. It is as yet unclear what type(s) of features can be bound to object indexicals, or that all types of features can be bound to object indexicals. It is also debated whether perceptual and conceptual features are bound to the same set of object indexicals or separate sets of object indexicals (see Murez, Smortchkova & Strickland, 2020; Brody, 2020; Yu & Lau, 2023).

2 Note that the exact duration of the NSW effect is under debate; some reports suggested that it could sustain for > 3 s (Ruchkin et al., 1992), while some other reports suggested that it terminates ∼ 1 s after memorandum onset (Fukuda, Mance & Vogel, 2015). The duration of the CDA is unclear as well (e.g., Perez & Vogel, 2012; Li & Noguchi, 2022). Here we simply note that the NSW and the CDA have broadly similar time courses in the sense that they are both late sustaining components.

3 In contrast to these studies on visual objects, the results of Morrison et al. (2020) suggest that holding 1, 3, 5 visually-presented *digits* does not drive a sustained parietal effect, at least from visual inspection of their figures, although the electrodes selected in that study (P3, Pz, P4) were less occipital compared to other NSW studies. Digits may encourage phonological rehearsal, whose load is manifested more in frontal and/or central electrodes (Alunni-Menichini et al., 2014; Lefebvre & Jolicœur, 2016; Simal & Jolicoeur, 2020; Bidelman, Brown & Bashivan, 2021).

4 Note that there have been many fMRI studies comparing events (using pictorial or sentential stimuli) against various types of baselines (e.g., Centelles et al., 2011; Quadflieg, Gentile & Rossion, 2015; Isik et al., 2017; Okruszek et al., 2017; Matchin et al., 2019b), but due to the relatively poor temporal resolution of fMRI, it is hard to disentangle the extraction and maintenance phases, while the maintenance phase is of our current interest.

5 Other possibilities will be discussed in Section 3.2.

6 Notably, the reaction times seemed to be longer for Experiment 2 compared to Experiment 1, despite similar stimulus-onset asynchrony across the picture and the linguistic probe. This is in line with the observation in J. Zhang et al. (2021): although holding content in the activity-silent mode for a longer time does not always come with a lower behavioral accuracy, it can come with a longer reaction time to the probe.

## References

1. Altmann, G., & Ekves, Z. (2019). Events as intersecting object histories: A new theory of event representation. Psychological Review, 126(6), 817.

2. Altmann, L. J., Saleem, A., Kendall, D., Heilman, K. M., & Rothi, L. J. G. (2006). Orthographic directionality and thematic role illustration in English and Arabic. Brain and Language, 97(3), 306–316.

3. Alunni-Menichini, K., Guimond, S., Bermudez, P., Nolden, S., Lefebvre, C., & Jolicoeur, P. (2014). Saturation of auditory short-term memory causes a plateau in the sustained anterior negativity event-related potential. Brain Research, 1592, 55–64.

4. Anderson, J., & Hastie, R. (1974). Individuation and reference in memory: Proper names and definite descriptions. Cognitive Psychology, 6(4), 495–514.

5. Awh, E., Barton, B., & Vogel, E. K. (2007). Visual working memory represents a fixed number of items regardless of complexity. Psychological Science, 18(7), 622–628.

6. Balaban, H., & Luria, R. (2016). Object representations in visual working memory change according to the task context. Cortex, 81, 1–13.

7. Baldassano, C., Chen, J., Zadbood, A., Pillow, J. W., Hasson, U., & Norman, K. A. (2017). Discovering event structure in continuous narrative perception and memory. Neuron, 95(3), 709–721.

8. Balestrieri, E., Ronconi, L., & Melcher, D. (2022). Shared resources between visual attention and visual working memory are allocated through rhythmic sampling. European Journal of Neuroscience, 55(11-12), 3040–3053.

9. Barrett, A. M., Kim, M., Crucian, G. P., & Heilman, K. M. (2002). Spatial bias: Effects of early reading direction on Korean subjects. Neuropsychologia, 40(7), 1003–1012.

10. Becke, A., Müller, N., Vellage, A., Schoenfeld, M. A., & Hopf, J. M. (2015). Neural sources of visual working memory maintenance in human parietal and ventral extrastriate visual cortex. NeuroImage, 110, 78–86.

11. Berggren, N., & Eimer, M. (2016). Does contralateral delay activity reflect working memory storage or the current focus of spatial attention within visual working memory?. Journal of Cognitive Neuroscience, 28(12), 2003–2020.

12. Bettencourt, K. C., & Xu, Y. (2016). Decoding the content of visual short-term memory under distraction in occipital and parietal areas. Nature Neuroscience, 19(1), 150–157.

13. Beukers, A. O., Buschman, T. J., Cohen, J. D., & Norman, K. A. (2021). Is activity silent working memory simply episodic memory?. Trends in Cognitive Sciences, 25(4), 284–293.

14. Bidelman, G. M., Brown, J. A., & Bashivan, P. (2021). Auditory cortex supports verbal working memory capacity. NeuroReport, 32(2), 163.

15. Brainard, D. H. (1997). The Psychophysics Toolbox. Spatial Vision, 10(4), 433–436.

16. Brody, G. (2020). Indexing objects in vision and communication [Doctoral dissertation]. Central European University.

17. Bueti, D., & Walsh, V. (2009). The parietal cortex and the representation of time, space, number and other magnitudes. Philosophical Transactions of the Royal Society B: Biological Sciences, 364(1525), 1831–1840.

18. Cai, Y., Fulvio, J. M., Samaha, J., & Postle, B. R. (2022). Context-binding in visual working memory is reflected in bilateral event-related potentials, but not in contralateral delay activity. eNeuro, 9(6), 1–15.

19. Carey, S. (2009). The origin of concepts. Oxford University Press.

20. Centelles, L., Assaiante, C., Nazarian, B., Anton, J. L., & Schmitz, C. (2011). Recruitment of both the mirror and the mentalizing networks when observing social interactions depicted by point-lights: a neuroimaging study. PLoS One, 6(1), e15749.

21. Chatterjee, A., Maher, L. M., & Heilman, K. M. (1995). Spatial characteristics of thematic role representation. Neuropsychologia, 33(5), 643–648.

22. Chota, S., Leto, C., van Zantwijk, L., & Van der Stigchel, S. (2022). Attention rhythmically samples multi-feature objects in working memory. Scientific Reports, 12(1), 1–12.

23. Clevenger, P. E., & Hummel, J. E. (2014). Working memory for relations among objects. Attention, Perception, & Psychophysics, 76(7), 1933–1953.

24. Cohn, N., Paczynski, M., & Kutas, M. (2017). Not so secret agents: Event-related potentials to semantic roles in visual event comprehension. Brain and Cognition, 119, 1–9.

25. Coslett, H. B., & Saffran, E. (1991). Simultanagnosia: To see but not two see. Brain, 114(4), 1523–1545.

26. Cowan, N. (2001). The magical number 4 in short-term memory: A reconsideration of mental storage capacity. Behavioral and Brain Sciences, 24(1), 87–114.

27. Cruz Heredia, A. A., Dickerson, B., & Lau, E. (2022). Towards understanding sustained neural activity across syntactic dependencies. Neurobiology of Language, 3(1), 87–108.

28. de la Rosa, S., Choudhery, R. N., Curio, C., Ullman, S., Assif, L., & Buelthoff, H. H. (2014). Visual categorization of social interactions. Visual Cognition, 22(9-10), 1233–1271.

29. Delorme, A., & Makeig, S. (2004). EEGLAB: an open source toolbox for analysis of single-trial EEG dynamics including independent component analysis. Journal of Neuroscience Methods, 134(1), 9–21.

30. Diaz, G. K., Vogel, E. K., & Awh, E. (2021). Perceptual grouping reveals distinct roles for sustained slow wave activity and alpha oscillations in working memory. Journal of Cognitive Neuroscience, 33(7), 1354–1364.

31. Ding, X., Gao, Z., & Shen, M. (2017). Two equals one: two human actions during social interaction are grouped as one unit in working memory. Psychological Science, 28(9), 1311–1320.

32. Dobel, C., Diesendruck, G., & Bölte, J. (2007). How writing system and age influence spatial representations of actions: A developmental, cross-linguistic study. Psychological Science, 18(6), 487–491.

33. Dobel, C., Enriquez-Geppert, S., Zwitserlood, P., & Bölte, J. (2014). Literacy shapes thought: The case of event representation in different cultures. Frontiers in Psychology, 5, 290.

34. Dobel, C., Gumnior, H., Bölte, J., & Zwitserlood, P. (2007). Describing scenes hardly seen. Acta Psychologica, 125(2), 129–143.

35. Dowty, D. (1991). Thematic proto-roles and argument selection. Language, 67(3), 547–619.

36. Duncan, D., van Moorselaar, D., & Theeuwes, J. (2023). Pinging the brain to reveal the hidden attentional priority map. bioRxiv.

37. Eichenbaum, H., & Cohen, N. J. (2014). Can we reconcile the declarative memory and spatial navigation views on hippocampal function?. Neuron, 83(4), 764–770.

38. Fan, Y., & Luo, H. (2022). Reactivating ordinal position information from auditory sequence memory in human brains. bioRxiv.

39. Fan, Y., Han, Q., Guo, S., & Luo, H. (2021). Distinct neural representations of content and ordinal structure in auditory sequence memory. Journal of Neuroscience, 41(29), 6290–6303.

40. Fan, Z., Muthukumaraswamy, S. D., Singh, K. D., & Shapiro, K. (2012). The role of sustained posterior brain activity in the serial chaining of two cognitive operations: A MEG study. Psychophysiology, 49(8), 1133–1144.

41. Feldmann-Wüstefeld, T. (2021). Neural measures of working memory in a bilateral change detection task. Psychophysiology, 58(1), e13683.

42. Feldmann-Wüstefeld, T., Vogel, E. K., & Awh, E. (2018). Contralateral delay activity indexes working memory storage, not the current focus of spatial attention. Journal of Cognitive Neuroscience, 30(8), 1185–1196.

43. Fitch, W. T. (2020). Animal cognition and the evolution of human language: why we cannot focus solely on communication. Philosophical Transactions of the Royal Society B, 375(1789), 20190046.

44. Fornaciai, M., & Park, J. (2020). Neural dynamics of serial dependence in numerosity perception. Journal of Cognitive Neuroscience, 32(1), 141–154.

45. Frankland, S. M., & Greene, J. D. (2015). An architecture for encoding sentence meaning in left mid-superior temporal cortex. Proceedings of the National Academy of Sciences, 112(37), 11732–11737.

46. Frankland, S. M., & Greene, J. D. (2020). Two ways to build a thought: distinct forms of compositional semantic representation across brain regions. Cerebral Cortex, 30(6), 3838–3855.

47. Friedman-Hill, S. R., Robertson, L. C., & Treisman, A. (1995). Parietal contributions to visual feature binding: evidence from a patient with bilateral lesions. Science, 269(5225), 853–855.

48. Fukuda, K., Mance, I., & Vogel, E. K. (2015). α power modulation and event-related slow wave provide dissociable correlates of visual working memory. Journal of Neuroscience, 35(41), 14009–14016.

49. Gao, Z., Ding, X., Yang, T., Liang, J., & Shui, R. (2013). Coarse-to-fine construction for high-resolution representation in visual working memory. PLoS One, 8(2), e57913.

50. Gao, Z., Ding, X., Yang, T., Liang, J., & Shui, R. (2013). Coarse-to-fine construction for high-resolution representation in visual working memory. PLoS One, 8(2), e57913.

51. Gaston, P. E. (2020). The role of syntactic prediction in auditory word recognition (Doctoral dissertation, University of Maryland, College Park).

52. Geminiani, G., Bisiach, E., Berti, A., & Rusconi, M. L. (1995). Analogical representation and language structure. Neuropsychologia, 33(11), 1565–1574.

53. Gerber, E. M. (2021). permutest (https://www.mathworks.com/matlabcentral/fileexchange/71737-permutest), MATLAB Central File Exchange. Retrieved December 29, 2021.

54. Hafri, A., Papafragou, A., & Trueswell, J. C. (2013). Getting the gist of events: recognition of two-participant actions from brief displays. Journal of Experimental Psychology: General, 142(3), 880.

55. Hafri, A., Trueswell, J. C., & Epstein, R. A. (2017). Neural representations of observed actions generalize across static and dynamic visual input. Journal of Neuroscience, 37(11), 3056–3071.

56. Hakim, N., Adam, K. C., Gunseli, E., Awh, E., & Vogel, E. K. (2019). Dissecting the neural focus of attention reveals distinct processes for spatial attention and object-based storage in visual working memory. Psychological Science, 30(4), 526–540.

57. Huang, Q., Zhang, H., & Luo, H. (2021). Sequence structure organizes items in varied latent states of working memory neural network. eLife, 10, e67589.

58. Hultén, A., Karvonen, L., Laine, M., & Salmelin, R. (2014). Producing speech with a newly learned morphosyntax and vocabulary: an magnetoencephalography study. Journal of Cognitive Neuroscience, 26(8), 1721–1735.

59. Hummel, J. E., & Holyoak, K. J. (2003). A symbolic-connectionist theory of relational inference and generalization. Psychological Review, 110(2), 220.

60. Hyun, J. S., Woodman, G. F., Vogel, E. K., Hollingworth, A., & Luck, S. J. (2009). The comparison of visual working memory representations with perceptual inputs. Journal of Experimental Psychology: Human Perception and Performance, 35(4), 1140.

61. Isik, L., Koldewyn, K., Beeler, D., & Kanwisher, N. (2017). Perceiving social interactions in the posterior superior temporal sulcus. Proceedings of the National Academy of Sciences, 114(43), E9145–E9152.

62. Ivanova, A. A., Mineroff, Z., Zimmerer, V., Kanwisher, N., Varley, R., & Fedorenko, E. (2021). The language network is recruited but not required for nonverbal event semantics. Neurobiology of Language, 2(2), 176–201.

63. Jackendoff, R. (1987). On beyond zebra: The relation of linguistic and visual information. Cognition, 26(2), 89–114.

64. JASP Team (2022). JASP (Version 0.16.1) [Computer software].

65. Jessop, A., & Chang, F. (2020). Thematic role information is maintained in the visual object-tracking system. Quarterly Journal of Experimental Psychology, 73(1), 146–163.

66. Jessop, A., & Chang, F. (2022). Thematic role tracking difficulties across multiple visual events influences role use in language production. Visual Cognition, 30(3), 151–173.

67. Jia, J., Liu, L., Fang, F., & Luo, H. (2017). Sequential sampling of visual objects during sustained attention. PLoS Biology, 15(6), e2001903.

68. Jia, K., Li, Y., Gong, M., Huang, H., Wang, Y., & Li, S. (2021). Perceptual learning beyond perception: mnemonic representation in early visual cortex and intraparietal sulcus. Journal of Neuroscience, 41(20), 4476–4486.

69. Johnson, M. G. (1967). Syntatic position and rated meaning. Journal of Verbal Learning and Verbal Behavior, 6(2), 240–246.

70. Jouen, A. L., Ellmore, T. M., Madden, C. J., Pallier, C., Dominey, P. F., & Ventre-Dominey, J. (2015). Beyond the word and image: characteristics of a common meaning system for language and vision revealed by functional and structural imaging. NeuroImage, 106, 72–85.

71. Kaan, E., Harris, A., Gibson, E., & Holcomb, P. (2000). The P600 as an index of syntactic integration difficulty. Language and Cognitive Processes, 15(2), 159–201.

72. Kahneman, D., Treisman, A., & Gibbs, B. J. (1992). The reviewing of object files: Object-specific integration of information. Cognitive Psychology, 24(2), 175–219.

73. Kako, E. (2006a). Thematic role properties of subjects and objects. Cognition, 101(1), 1–42.

74. Kako, E. (2006b). The semantics of syntactic frames. Language and Cognitive Processes, 21(5), 562–575.

75. Kamiński, J., & Rutishauser, U. (2020). Between persistently active and activity-silent frameworks: novel vistas on the cellular basis of working memory. Annals of the New York Academy of Sciences, 1464(1), 64–75.

76. Kazandjian, S., Gaash, E., Love, I. Y., Zivotofsky, A. Z., & Chokron, S. (2011). Spatial representation of action phrases among bidirectional readers: The effect of language environment and sentence complexity. Social Psychology, 42(3), 249.

77. Klaver, P., Smid, H. G., & Heinze, H. J. (1999). Representations in human visual short-term memory: an event-related brain potential study. Neuroscience Letters, 268(2), 65–68.

78. Kleiner, M., Brainard, D., & Pelli, D. (2007). What’s new in Psychtoolbox-3? Perception, 36, 14.

79. Knops, A., Piazza, M., Sengupta, R., Eger, E., & Melcher, D. (2014). A shared, flexible neural map architecture reflects capacity limits in both visual short-term memory and enumeration. Journal of Neuroscience, 34(30), 9857–9866.

80. Konkel, A., & Cohen, N. J. (2009). Relational memory and the hippocampus: representations and methods. Frontiers in Neuroscience, 23.

81. Kreither, J., Papaioannou, O., & Luck, S. J. (2022). Active working memory and simple cognitive operations. Journal of Cognitive Neuroscience, 34(2), 313–331.

82. Lalisse, M., & Smolensky, P. (2021). Distributed neural encoding of binding to thematic roles. arXiv preprint arXiv:2110.12342.

83. Lapinskaya, N., Uzomah, U., Bedny, M., & Lau, E. (2016). Electrophysiological signatures of event words: dissociating syntactic and semantic category effects in lexical processing. Neuropsychologia, 93, 151–157.

84. Lau, E. (2018). Neural indices of structured sentence representation: State of the art. In Psychology of learning and motivation (pp. 117–142). Elsevier.

85. Lau, E., & Liao, C. H. (2018). Linguistic structure across time: ERP responses to coordinated and uncoordinated noun phrases. Language, Cognition and Neuroscience, 33(5), 633–647.

86. Lau, E., & Yang, X. (2023). *Neural response to coordination reflects interpretive processes, not syntactic connectedness [conference presentation]*. Human Sentence Processing 2023, Pittsburgh, USA.

87. Leavitt, M. L., Mendoza-Halliday, D., & Martinez-Trujillo, J. C. (2017). Sustained activity encoding working memories: not fully distributed. Trends in Neurosciences, 40(6), 328–346.

88. Lefebvre, C., & Jolicœur, P. (2016). Memory for pure tone sequences without contour. Brain Research, 1640, 222–231.

89. Leslie, A. M., Xu, F., Tremoulet, P. D., & Scholl, B. J. (1998). Indexing and the object concept: developing ’what’ and ‘where’ systems. Trends in Cognitive Sciences, 2(1), 10–18.

90. Lewis-Peacock, J. A., Drysdale, A. T., Oberauer, K., & Postle, B. R. (2012). Neural evidence for a distinction between short-term memory and the focus of attention. Journal of Cognitive Neuroscience, 24(1), 61–79.

91. Li, Y., & Noguchi, Y. (2022). Neural correlates of a load-dependent decline in visual working memory. Cerebral Cortex Communications, 3(2), tgac015.

92. Liu, C., Wang, R., Li, L., Ding, G., Yang, J., & Li, P. (2020). Effects of encoding modes on memory of naturalistic events. Journal of Neurolinguistics, 53, 100863.

93. Lo, C. W., & Brennan, J. R. (2021). EEG Correlates of Long-Distance Dependency Formation in Mandarin Wh-Questions. Frontiers in Human Neuroscience, 15, 591613.

94. Lopez-Calderon, J., & Luck, S. J. (2014). ERPLAB: an open-source toolbox for the analysis of event-related potentials. Frontiers in Human Neuroscience, 8, 213.

95. Lu, X., Dai, A., Guo, Y., Shen, M., & Gao, Z. (2022). Is the social chunking of agent actions in working memory resource-demanding?. Cognition, 229, 105249.

96. Lu, X., Huang, J., Yi, Y., Shen, M., Weng, X., & Gao, Z. (2016). Holding biological motion in working memory: An fMRI study. Frontiers in Human Neuroscience, 10, 251.

97. Luck, S. J., & Vogel, E. K. (1997). The capacity of visual working memory for features and conjunctions. Nature, 390(6657), 279–281.

98. Luck, S. J., & Vogel, E. K. (2013). Visual working memory capacity: from psychophysics and neurobiology to individual differences. Trends in Cognitive Sciences, 17(8), 391–400.

99. Lundqvist, M., Rose, J., Warden, M. R., Buschman, T., Miller, E. K., & Herman, P. (2021). A Hot-Coal theory of Working Memory. bioRxiv.

100. Luria, R., Balaban, H., Awh, E., & Vogel, E. K. (2016). The contralateral delay activity as a neural measure of visual working memory. Neuroscience & Biobehavioral Reviews, 62, 100–108.

101. Luyckx, F., Nili, H., Spitzer, B., & Summerfield, C. (2019). Neural structure mapping in human probabilistic reward learning. eLife, 8, e42816.

102. Ma, F., Xu, J., Li, X., Wang, P., Wang, B., & Liu, B. (2018). Investigating the neural basis of basic human movement perception using multi-voxel pattern analysis. Experimental Brain Research, 236(3), 907–918.

103. Maass, A., & Russo, A. (2003). Directional bias in the mental representation of spatial events: Nature or culture?. Psychological Science, 14(4), 296–301.

104. Madnani, N., Boyd-Graber, J., & Resnik, P. (2010). Measuring transitivity using untrained annotators. In Proceedings of the NAACL HLT 2010 workshop on creating speech and language data with Amazon’s Mechanical Turk (pp. 188–194).

105. Mahon, B. Z., & Kemmerer, D. (2020). Interactions between language, thought, and perception: Cognitive and neural perspectives. Cognitive Neuropsychology, 37(5-6), 235–240.

106. Maris, E., & Oostenveld, R. (2007). Nonparametric statistical testing of EEG-and MEG-data. Journal of Neuroscience Methods, 164(1), 177–190.

107. Marr, D. (1976). Early processing of visual information. Philosophical Transactions of the Royal Society of London. B, Biological Sciences, 275(942), 483–519.

108. Martin, A. E., & Doumas, L. A. (2017). A mechanism for the cortical computation of hierarchical linguistic structure. PLoS Biology, 15(3), e2000663.

109. Martin, B., Wiener, M., & van Wassenhove, V. (2017). A Bayesian perspective on accumulation in the magnitude system. Scientific reports, 7(1), 1–14.

110. Matchin, W., Brodbeck, C., Hammerly, C., & Lau, E. (2019a). The temporal dynamics of structure and content in sentence comprehension: Evidence from fMRI-constrained MEG. Human Brain Mapping, 40(2), 663–678.

111. Matchin, W., Liao, C. H., Gaston, P., & Lau, E. (2019b). Same words, different structures: An fMRI investigation of argument relations and the angular gyrus. Neuropsychologia, 125, 116–128.

112. Matsuyoshi, D., Ikeda, T., Sawamoto, N., Kakigi, R., Fukuyama, H., & Osaka, N. (2010). Task-irrelevant memory load induces inattentional blindness without temporo-parietal suppression. Neuropsychologia, 48(10), 3094–3101.

113. McCants, C. W., Katus, T., & Eimer, M. (2020). Task goals modulate the activation of part-based versus object-based representations in visual working memory. Cognitive Neuroscience, 11(1-2), 92–100.

114. McKinnon, R., & Osterhout, L. (1996). Constraints on movement phenomena in sentence processing: Evidence from event-related brain potentials. Language and cognitive processes, 11(5), 495–524.

115. Mecklinger, A., & Pfeifer, E. (1996). Event-related potentials reveal topographical and temporal distinct neuronal activation patterns for spatial and object working memory. Cognitive Brain Research, 4(3), 211–224.

116. Menenti, L., Segaert, K., & Hagoort, P. (2012). The neuronal infrastructure of speaking. Brain and Language, 122(2), 71–80.

117. Mitchell, D. J., & Cusack, R. (2008). Flexible, capacity-limited activity of posterior parietal cortex in perceptual as well as visual short-term memory tasks. Cerebral Cortex, 18(8), 1788–1798.

118. Mitchell, D. J., & Cusack, R. (2011). The temporal evolution of electromagnetic markers sensitive to the capacity limits of visual short-term memory. Frontiers in Human Neuroscience, 5, 18.

119. Morrison, C., Kamal, F., Le, K., & Taler, V. (2020). Monolinguals and bilinguals respond differently to a delayed matching-to-sample task: An ERP study. Bilingualism: Language and Cognition, 23(4), 858–868.

120. Murez, M., Smortchkova, J., & Strickland, B. (2020). The Mental Files Theory of Singular Thought. In Singular Thought and Mental Files (pp.107–142).

121. Naughtin, C. K., Mattingley, J. B., & Dux, P. E. (2016). Distributed and overlapping neural substrates for object individuation and identification in visual short-term memory. Cerebral Cortex, 26(2), 566–575.

122. Ngiam, W. X., Foster, J. J., Adam, K. C., & Awh, E. (2022). Distinguishing guesses from fuzzy memories: Further evidence for item limits in visual working memory. Attention, Perception, & Psychophysics.

123. O’Reilly, R. C., Ranganath, C., & Russin, J. L. (2022). The Structure of Systematicity in the Brain. Current Directions in Psychological Science, 31(2), 124–130.

124. O’Connell, R. G., & Kelly, S. P. (2021). Neurophysiology of human perceptual decision-making. Annual Review of Neuroscience, 44, 495–516.

125. O’connell, R. G., Dockree, P. M., & Kelly, S. P. (2012). A supramodal accumulation-to-bound signal that determines perceptual decisions in humans. Nature Neuroscience, 15(12), 1729–1735.

126. Okruszek, Ł., Wordecha, M., Jarkiewicz, M., Kossowski, B., Lee, J., & Marchewka, A. (2018). Brain correlates of recognition of communicative interactions from biological motion in schizophrenia. Psychological Medicine, 48(11), 1862–1871.

127. Oldfield, R. C. (1971). The assessment and analysis of handedness: the Edinburgh inventory. Neuropsychologia, 9(1), 97–113.

128. Paparella, I., & Papeo, L. (2022). Chunking by social relationship in working memory. Visual Cognition, 1–17.

129. Park, H. B., Zhang, W., & Hyun, J. S. (2017). Dissociating models of visual working memory by reaction-time distribution analysis. Acta Psychologica, 173, 21–31.

130. Peng, Y., Javangula, P. R., Lu, H., & Holyoak, K. J. (2018). Behavioral oscillations in verification of relational role bindings. In C. Kalish, M. Rau, J. Zhu & T. T. Rogers (Eds.), Proceedings of the 40th Annual Conference of the Cognitive Science Society. Austin, TX: Cognitive Science Society.

131. Perez, V. B., & Vogel, E. K. (2012). What ERPs can tell us about working memory. In The Oxford handbook of event-related potential components (pp. 361–372).

132. Peters, B., Kaiser, J., Rahm, B., & Bledowski, C. (2021). Object-based attention prioritizes working memory contents at a theta rhythm. Journal of Experimental Psychology: General, 150(6), 1250.

133. Peterson, D. J., Gözenman, F., Arciniega, H., & Berryhill, M. E. (2015). Contralateral delay activity tracks the influence of Gestalt grouping principles on active visual working memory representations. Attention, Perception, & Psychophysics, 77(7), 2270–2283.

134. Pion-Tonachini, L., Kreutz-Delgado, K., & Makeig, S. (2019). ICLabel: An automated electroencephalographic independent component classifier, dataset, and website. NeuroImage, 198, 181–197.

135. Pitcher, D., & Ungerleider, L. G. (2021). Evidence for a third visual pathway specialized for social perception. Trends in Cognitive Sciences, 25(2), 100–110.

136. Pomper, U., & Ansorge, U. (2021). Theta-rhythmic oscillation of working memory performance. Psychological Science, 32(11), 1801–1810.

137. Pylyshyn, Z. (1989). The role of location indexes in spatial perception: A sketch of the FINST spatial-index model. Cognition, 32(1), 65–97.

138. Pylyshyn, Z. W. (2001). Visual indexes, preconceptual objects, and situated vision. Cognition, 80(1-2), 127–158.

139. Quadflieg, S., & Koldewyn, K. (2017). The neuroscience of people watching: how the human brain makes sense of other people’s encounters. Annals of the New York Academy of Sciences, 1396(1), 166–182.

140. Quilty-Dunn, J., & Green, E. J. (2021). Perceptual attribution and perceptual reference. Philosophy and Phenomenological Research.

141. Quilty-Dunn, J., Porot, N., & Mandelbaum, E. (2022). The best game in town: The re-emergence of the language of thought hypothesis across the cognitive sciences. Behavioral and Brain Sciences.

142. Rabbitt, L. R., Roberts, D. M., McDonald, C. G., & Peterson, M. S. (2017). Neural activity reveals perceptual grouping in working memory. International Journal of Psychophysiology, 113, 40–45.

143. Rafal, R. (2001). Balint’s syndrome. In Handbook of neuropsychology (pp. 121–141).

144. Reisinger, D., Rudinger, R., Ferraro, F., Harman, C., Rawlins, K., & Van Durme, B. (2015). Semantic proto-roles. Transactions of the Association for Computational Linguistics, 3, 475–488.

145. Rissman, L., & Lupyan, G. (2022). A dissociation between conceptual prominence and explicit category learning: Evidence from agent and patient event roles. Journal of Experimental Psychology: General, 151(7), 1707.

146. Rissman, L., & Majid, A. (2019). Thematic roles: Core knowledge or linguistic construct?. Psychonomic Bulletin & Review, 26(6), 1850–1869.

147. Robitaille, N., Marois, R., Todd, J., Grimault, S., Cheyne, D., & Jolicœur, P. (2010). Distinguishing between lateralized and nonlateralized brain activity associated with visual short-term memory: fMRI, MEG, and EEG evidence from the same observers. NeuroImage, 53(4), 1334–1345.

148. Rose, N. S., LaRocque, J. J., Riggall, A. C., Gosseries, O., Starrett, M. J., Meyering, E. E., & Postle, B. R. (2016). Reactivation of latent working memories with transcranial magnetic stimulation. Science, 354(6316), 1136–1139.

149. Ruchkin, D. S., Johnson Jr, R., Grafman, J., Canoune, H., & Ritter, W. (1992). Distinctions and similarities among working memory processes: An event-related potential study. Cognitive Brain Research, 1(1), 53–66.

150. Shastri, L. (1999). Advances in Shruti—A neurally motivated model of relational knowledge representation and rapid inference using temporal synchrony. Applied Intelligence, 11(1), 79–108.

151. Shastri, L. (2002). Episodic memory and cortico–hippocampal interactions. Trends in Cognitive Sciences, 6(4), 162–168.

152. Shastri, L. (2007). SHRUTI: A neurally motivated architecture for rapid, scalable inference. In Perspectives of Neural-Symbolic Integration (pp. 183–203). Springer, Berlin, Heidelberg.

153. Shastri, L., & Ajjanagadde, V. (1993). From simple associations to systematic reasoning: A connectionist representation of rules, variables and dynamic bindings using temporal synchrony. Behavioral and Brain Sciences, 16(3), 417–451.

154. Shen, M., Chen, J., Yang, X., Dong, H., Chen, H., & Zhou, J. (2021). The storage mechanism of dynamic relations in visual working memory. Cognition, 209, 104571.

155. Simal, A., & Jolicoeur, P. (2020). Scanning acoustic short-term memory: Evidence for two subsystems with different time-course and memory strength. International Journal of Psychophysiology, 155, 105–117.

156. Song, J. H., & Jiang, Y. (2006). Visual working memory for simple and complex features: An fMRI study. NeuroImage, 30(3), 963–972.

157. Spitzer, B., Waschke, L., & Summerfield, C. (2017). Selective overweighting of larger magnitudes during noisy numerical comparison. Nature Human Behaviour, 1(8), 1–8.

158. Sprouse, J., Kucerova, I., Park, J., Cerrone, P., Schrum, N., & Lau, E. F. (in preparation). Sustained anterior negativities and movement dependencies in syntax. Department of Linguistics, University of Connecticut.

159. Stahl, A. E., & Feigenson, L. (2014). Social knowledge facilitates chunking in infancy. Child Development, 85(4), 1477–1490.

160. Stokes, M. G. (2015). ‘Activity-silent’ working memory in prefrontal cortex: a dynamic coding framework. Trends in Cognitive Sciences, 19(7), 394–405.

161. Strickland, B. (2017). Language reflects “core” cognition: A new theory about the origin of cross-linguistic regularities. Cognitive Science, 41(1), 70–101.

162. Summerfield, C., Luyckx, F., & Sheahan, H. (2020). Structure learning and the posterior parietal cortex. Progress in Neurobiology, 184, 101717.

163. Ten Oever, S., De Weerd, P., & Sack, A. T. (2020). Phase-dependent amplification of working memory content and performance. Nature Communications, 11(1), 1–8.

164. Thompson, C. K., Bonakdarpour, B., Fix, S. C., Blumenfeld, H. K., Parrish, T. B., Gitelman, D. R., & Mesulam, M. M. (2007). Neural correlates of verb argument structure processing. Journal of Cognitive Neuroscience, 19(11), 1753–1767.

165. Thyer, W., Adam, K. C. S., Diaz, G. K., Velázquez Sánchez, I. N., Vogel, E. K., Awh, E. (2022). Storage in visual working memory recruits a content-independent pointer system. Psychological Science.

166. Todd, J. J., & Marois, R. (2004). Capacity limit of visual short-term memory in human posterior parietal cortex. Nature, 428(6984), 751–754.

167. Treisman, A. (1998). Feature binding, attention and object perception. Philosophical Transactions of the Royal Society of London. Series B: Biological Sciences, 353(1373), 1295–1306.

168. Ünal, E., Ji, Y., & Papafragou, A. (2021). From event representation to linguistic meaning. Topics in Cognitive Science, 13(1), 224–242.

169. Vogel, E. K., & Machizawa, M. G. (2004). Neural activity predicts individual differences in visual working memory capacity. Nature, 428(6984), 748–751.

170. Walbrin, J., & Koldewyn, K. (2019). Dyadic interaction processing in the posterior temporal cortex. NeuroImage, 198, 296–302.

171. Walbrin, J., Downing, P., & Koldewyn, K. (2018). Neural responses to visually observed social interactions. Neuropsychologia, 112, 31–39.

172. Walsh, V. (2003). A theory of magnitude: common cortical metrics of time, space and quantity. Trends in Cognitive Sciences, 7(11), 483–488.

173. Wang, J., Cherkassky, V. L., Yang, Y., Chang, K. M. K., Vargas, R., Diana, N., & Just, M. A. (2016). Identifying thematic roles from neural representations measured by functional magnetic resonance imaging. Cognitive Neuropsychology, 33(3-4), 257–264.

174. Wang, X. J. (2021). 50 years of mnemonic persistent activity: quo vadis?. Trends in Neurosciences, 44(11), 888–902.

175. Williams, A. (2015). Arguments in syntax and semantics. Cambridge University Press.

176. Williams, A. (2021). Events in semantics. In The Cambridge Handbook of the Philosophy of Language (pp. 366–386).

177. Williams, A., Reddigari, S., & Pylkkänen, L. (2017). Early sensitivity of left perisylvian cortex to relationality in nouns and verbs. Neuropsychologia, 100, 131–143.

178. Wilson, V. A., Zuberbühler, K., & Bickel, B. (2022). The evolutionary origins of syntax: Event cognition in nonhuman primates. Science Advances, 8(25), eabn8464.

179. Wolff, M. J., Ding, J., Myers, N. E., & Stokes, M. G. (2015). Revealing hidden states in visual working memory using electroencephalography. Frontiers in Systems Neuroscience, 9, 123.

180. Wolff, M. J., Jochim, J., Akyürek, E. G., & Stokes, M. G. (2017). Dynamic hidden states underlying working-memory-guided behavior. Nature Neuroscience, 20(6), 864–871.

181. Wolff, M. J., Jochim, J., Akyürek, E. G., Buschman, T. J., & Stokes, M. G. (2020). Drifting codes within a stable coding scheme for working memory. PLoS Biology, 18(3), e3000625.

182. Wolff, M. J., Kandemir, G., Stokes, M. G., & Akyürek, E. G. (2019). Impulse responses reveal unimodal and bimodal access to visual and auditory working memory. bioRxiv, 623835.

183. Wurm, M. F., & Caramazza, A. (2019). Distinct roles of temporal and frontoparietal cortex in representing actions across vision and language. Nature Communications, 10(1), 1–10.

184. Wurm, M. F., & Lingnau, A. (2015). Decoding actions at different levels of abstraction. Journal of Neuroscience, 35(20), 7727–7735.

185. Wurm, M. F., Ariani, G., Greenlee, M. W., & Lingnau, A. (2016). Decoding concrete and abstract action representations during explicit and implicit conceptual processing. Cerebral Cortex, 26(8), 3390–3401.

186. Wurm, M. F., Caramazza, A., & Lingnau, A. (2017). Action categories in lateral occipitotemporal cortex are organized along sociality and transitivity. Journal of Neuroscience, 37(3), 562–575.

187. Wutz, A., Zazio, A., & Weisz, N. (2020). Oscillatory bursts in parietal cortex reflect dynamic attention between multiple objects and ensembles. Journal of Neuroscience, 40(36), 6927–6937.

188. Xu, Y. (2007). The role of the superior intraparietal sulcus in supporting visual short-term memory for multifeature objects. Journal of Neuroscience, 27(43), 11676–11686.

189. Xu, Y. (2017). Reevaluating the sensory account of visual working memory storage. Trends in Cognitive Sciences, 21(10), 794–815.

190. Xu, Y., & Chun, M. M. (2006). Dissociable neural mechanisms supporting visual short-term memory for objects. Nature, 440(7080), 91–95.

191. Xu, Y., & Chun, M. M. (2007). Visual grouping in human parietal cortex. Proceedings of the National Academy of Sciences, 104(47), 18766–18771.

192. Xu, Y., & Chun, M. M. (2009). Selecting and perceiving multiple visual objects. Trends in Cognitive Sciences, 13(4), 167–174.

193. Yang, H., He, C., Han, Z., & Bi, Y. (2020). Domain-specific functional coupling between dorsal and ventral systems during action perception. Scientific Reports, 10(1), 1–14.

194. Yano, M., & Koizumi, M. (2018). Processing of non-canonical word orders in (in) felicitous contexts: Evidence from event-related brain potentials. Language, Cognition and Neuroscience, 33(10), 1340–1354.

195. Yano, M., & Koizumi, M. (2021). The role of discourse in long-distance dependency formation. Language, Cognition and Neuroscience, 36(6), 711–729.

196. Yeung, N., & Sanfey, A. G. (2004). Independent coding of reward magnitude and valence in the human brain. Journal of Neuroscience, 24(28), 6258–6264.

197. Yu, X., & Lau, E. (2023). The Binding Problem 2.0: Beyond Perceptual Features. Cognitive Science, 47(2), e13244.

198. Zadbood, A., Chen, J., Leong, Y. C., Norman, K. A., & Hasson, U. (2017). How we transmit memories to other brains: constructing shared neural representations via communication. Cerebral Cortex, 27(10), 4988–5000.

199. Zhang, J., Ye, C., Sun, H. J., Zhou, J., Liang, T., Li, Y., & Liu, Q. (2021). The passive state: A protective mechanism for information in working memory tasks. Journal of Experimental Psychology: Learning, Memory, and Cognition.

200. Zhang, W., & Luck, S. J. (2008). Discrete fixed-resolution representations in visual working memory. Nature, 453(7192), 233–235.

201. Zhou, H., Su, C., Wu, J., Li, J., Lu, X., Gong, L., … & Hu, Y. (2022). A domain-general frontoparietal network interacts with domain-preferential intermediate pathways to support working memory task. Cerebral Cortex.

202. Zhou, L., & Thomas, R. D. (2015). Principal component analysis of the memory load effect in a change detection task. Vision Research, 110, 1–6.

203. Zhu, S. D., Zhang, L. A., & von der Heydt, R. (2020). Searching for object pointers in the visual cortex. Journal of Neurophysiology, 123(5), 1979–1994.

